# Single-vessel transcriptome map pathological landscapes and reveal NR2F2-mediated smooth muscle cell phenotype acquisition in capillary malformations

**DOI:** 10.1101/2025.09.02.673874

**Authors:** Vi Nguyen, Irving Mao, Siwuxie He, Isabella Castellanos, Mackenzie Azuero, Marcelo L Hochman, Yin Rong, Laena Pernomian, Elliott H Chen, Harold I Friedman, Yan-Hua Chen, Qun Lu, Daping Fan, Camilla F Wenceslau, Dong-bao Chen, J Stuart Nelson, Anil G. Jegga, Yunguan Wang, Wenbin Tan

**Author notes:** Corresponding to: Yunguan Wang, Department of Pediatrics, University of Cincinnati College of Medicine, Cincinnati, Ohio 45229, USA;, Wenbin Tan, Department of Cell Biology and Anatomy, School of Medicine, and Department of Biomedical Engineering, College of Engineering and Computing, University of South Carolina, Columbia, South Carolina, 29209, USA. Equally contributes to this work (Vi Nguyen, Irving Mao).

## Abstract

**Background:** Capillary malformation (CM) is a congenital vascular anomaly affecting the skin, mucosa, and brain, yet the understanding of its vascular pathogenesis remains limited.

**Methods:** We applied spatial whole-transcriptome profiling (GeoMx) and gene set enrichment analysis within CM lesions at single vasculature level. Differentially expressed genes were validated by immunofluorescence staining. Phosphoproteomics was profiled to uncover lesion-wide phosphorylation sites on proteins. Single-cell RNA sequencing was performed on CM-derived induced pluripotent stem cells (iPSCs) to determine differentiation trajectories of lesional vascular lineages. In silico gene perturbation was used to predict candidate genes for modulating vascular pathological progression, followed by functional validation in CM iPSC-derived endothelial cells (ECs) using a Tet-on system.

**Results:** A spatial transcriptomic atlas was constructed, and pathological landscape of individual CM vasculature was delineated. CM vessels exhibited hallmarks of endothelial-to-mesenchymal transition (EndMT), including disruption of adherens junctions (AJs), vascular identity transitions, and metabolic remodeling. Phosphoproteomics confirmed that differentially phosphorylated proteins were enriched in EndMT- and AJ-related pathways. Aberrant expression of venous transcriptional factor NR2F2 was observed in lesional ECs and correlated with progressive enlargement from capillaries to larger-caliber vessels containing multiple layers of smooth muscle cells (SMCs). In CM iPSCs, differentiation course yielded reduced ECs but increased SMCs. *In silico* knockout simulation predicted NR2F2 as a crucial regulator of facilitating SMC phenotype in CM. Consistently, enforced NR2F2 expression during iPSC differentiation suppressed endothelial markers while inducing SMC-associated genes.

**Conclusions:** Single CM vasculature displays pathological hallmarks characterized by EndMT and AJ disruption, leading to progressive vascular remodeling. NR2F2 functions as a central regulatory factor orchestrating the acquisition of the SMC phenotype, thereby representing a potential therapeutic target in CM.

## Introduction

Congenital vascular malformations (VMs) are developmental disorders of the vasculature, affecting approximately 1.5% of the general population.^1^ According to ISSVA classification, congenital capillary malformations (CMs) include several subtypes: cutaneous and/or mucosal CM (also known as port-wine birthmarks, PWB), CM with neurological involvement (Sturge-Weber syndrome, SWS), diffuse CM with overgrowth, reticulate CM, and certain forms of telangiectasia.^2^ Among these, cutaneous CM is the most common, with a prevalence of 0.3%–0.9% in newborns.^3^ Notably, about 25% of infants with forehead CM lesions are at elevated risk of SWS, characterized by ipsilateral leptomeningeal angiomatosis.^4^ Clinical management of CMs remains challenging, often depending on lesion location and associated vascular anomalies. Pulsed dye laser (PDL) and photodynamic therapy (PDT) are standard treatments for cutaneous CM, showing comparable efficacy.^5,6^ However, complete clearance is achieved in fewer than 10% of patients,^7^ while approximately 20% of lesions are resistant to laser therapy.^8^ Despite three decades of clinical use, laser-based approaches have yielded only limited improvements in outcomes.^9,10^ A subset of CM endothelial cells (ECs) likely survive laser treatments, allowing the revascularization of lesional blood vessels after laser radiation.^11,12^ This clinical inadequacy demands an understanding of the pathological phenotypes of CMs for the development of paradigm-shifting modalities.

Histologically, CM lesions display diverse vascular abnormalities, including proliferation of ECs and smooth muscle cells (SMCs), replication of vascular basement membranes, disruption of vascular barriers, and progressive dilatation of vasculature.^12,13^ CM ECs exhibit stem-cell-like characteristics which have been posited as aberrant endothelial progenitor cells (EPCs),^12,14^ leading to impairments of EC differentiation.^12^ Several sporadic somatic mutations are shown to be related to vascular phenotypes of CMs, including the guanine nucleotide-binding protein, G alpha subunit q (GNAQ) (R183Q), G protein subunit alpha 11 (GNA11), phosphatidylinositol 3- kinase (PI3K), RAS P21 protein activator 1 (RASA1), STAM binding protein (STAMBP), and Ephrin type-B receptor 4.^2,15,16^ Cutaneous CM and CM/SWS vasculature have GNAQ (R183Q) allelic frequencies ranging from 1 to 12 percent (about 4.7% on average) in skin lesions.^15^ GNAQ (R183Q) is present in ECs or blood vessels,^11,17^ connective tissues, hair follicles, and glands in CM lesions, suggesting that GNAQ (R183Q) pluripotent cells may give rise to multiple lineages in cutaneous CM.^11^ EPCs with GNAQ (R183Q) can cause active phospholipase C (PLC) β3 signaling, increase angiopoietin-2 (ANGPT2), and produce enlarged capillary-like vessels.^18^ Intriguingly, GNAQ (R183Q) is embryonically lethal, and mutant ECs likely persist through mosaicism.^19^ The low allelic frequency and mosaic distribution of GNAQ (R183Q) suggest that additional factors contribute to CM pathology. Furthermore, patient-derived induced pluripotent stem cells (iPSCs) generated in our laboratory, while lacking GNAQ (R183Q), nonetheless recapitulate clinically relevant vascular phenotypes *in vitro* and *in vivo,*^20,21^ suggesting that epigenetic imprints within lesional cells may preserve disease phenotypes.

CM vasculature exhibits pronounced morphological and pathological heterogeneity, reflecting dynamic EC remodeling during disease progression. Increasing evidence points to a role for inflammation and metabolic reprogramming in CM pathology, as macrophages are frequently enriched in the perivascular environment of brain and skin lesions.^22,23^ However, the underlying molecular signatures remain largely unknown. To address this, we performed spatial whole- transcriptome atlas profiling of single CM vasculature. Our analysis delineated the pathological landscape of individual vasculatures with signatures of endothelial-to-mesenchymal transition (EndMT) and revealed aberrant expression of nuclear receptor subfamily 2 group F member 2 (NR2F2/COUP-TFII), coinciding with progression of vasculature enlargement. These findings were corroborated by CM phosphoproteomics, which demonstrated enrichment of EndMT and adherens junction (AJ)–related pathways among differentially phosphorylated proteins. Functionally, CM lesion–derived iPSCs showed impaired endothelial differentiation but an expansion of SMC lineages. Importantly, induction of NR2F2 expression during the endothelial progenitor stage promoted EndMT-associated SMC phenotypes in ECs.

## Methods

### Tissue preparation

This study (#1853132) was approved by the Institutional Review Board at Prisma Health Midlands. Some biopsy samples were reported in our previous studies.^11,12,24^ A total of 17 CM and 9 normal skin biopsy samples were used in this study, including (1) six surgically excised hypertrophic or nodular CM or biopsied CM/VM macular lesions and six de-identified surgically discarded normal skin tissues for GeoMx Spatial Whole Transcriptome Atlas (WTA) (Nanostring, Seattle, WA, USA), (2) six CM lesions and four normal skin controls for immunofluorescent (IF) staining assay to validate differentially expressed gene (DEG), and (3) five CM lesions and five normal skin controls for phosphoproteomics analysis. The biopsy tissues were 4% buffered formalin-fixed and parafilm-embedded (FFPE) for preparing sections (5 - 6 µm). The sample information was listed in supplementary table 1 Key Resource Table.

### Single vasculature Spatial WTA by GeoMx Digital Spatial Profiler

De-identified FFPE sections were stained with DAPI. The region of interest (ROI) containing individual blood vessel was picked. The sections were permeabilized and hybridized using RNA specific probes labeled with a unique barcoding system. Flow cells were assembled, followed by automation of multiple rounds of reporter probe binding and fluorescence imaging for barcode readouts. Each RNA appeared as a single bright spot in the sample and were digitally quantified in the image.

Differential expression analysis was performed with pseudobulk analysis (one vs. the rest) using DecoupleR (v1.8.0) and pyDEseq2 (v0.4.12).^25^ Absolute log fold change greater than 0.6 and FDR-adjusted P-value of 0.05 were used to select for differentially expressed genes (DEGs). Gene set enrichment analysis (GSEA) was performed with the package gseapy (v1.1.4 .) using the ‘KEGG’ and ‘Hallmark’ libraries from MsigDB.^26^

### CM iPSC culture and iEC differentiation

The generation of CM and normal iPSC lines were reported previously.^20,21^ Briefly, normal and CM iPSC lines were seeded into Geltrex-coated 6 well plate in mTeSR plus medium (STEMCELL, Cambridge, MA, USA) with 10µM Y27632. For iEC differentiation, the protocol was used in favor to arterial-like lineage commitment.^27–30^ The details were described in supplementary method. For iPSC lines with integration of Tet-on system for inducible expression of NR2F2, the iPSCs were electroporated using pPB-cT3G-cERP2-NR2F2^31^ (Addgene #192900, Watertown, MA, USA) plus a Super piggyBac Transposase expression vector (SBI#, PB210PA-1, Palo Alto, CA, USA). The cells were selected under 1.0-1.2 µg/ml puromycin to obtain the iPSC lines with a stable integration of Tet-on system. The NR2F2 expression was induced using doxycycline (Dox, 1 µg/ml) during the designated phase.

### Single-cell RNA sequencing (scRNA-seq)

The samples were collected for scRNA-seq at the following iPSC differentiation stages: iPSC (day 0), mesodermal cells (MC, day 4), progenitor cell mixture including EPCs and smooth muscle progenitor cells (SMPC) (day 8), and iECs (day 14). Parse Evercode™ single-cell library kit was used for library construction (Pars Bioscience, Seattle, WA, USA). Samples with biological duplicates per group were processed. Paired-end sequencing was performed on a NovaSeq 6000 system (Illumina, San Diego, CA, USA) for 50 K reads per cell. GSEA based pathway analysis was performed using the python package gesapy (v 1.1.4). The ‘prerank’ function was used to identify enriched pathways from DEG ranked by log foldchange. Pseudotime analysis was conducted using Diffusion Pseudotime (DPT, included in the scanpy package), which leverages the diffusion map framework to order cells along a continuum based on their transcriptional similarity.

### Bulk ATAC-seq and regulatory gene network (RGN) construction

Bulk ATAC-seq was performed on iECs using a kit from Diagenode (Denville, NJ, USA) with detailed description in supplementary method. Bowtie2^32^ was used for ATAC-seq alignment and MACS2^33^ for peak calling. The ATAC-seq heatmap was generated using Galaxy platform.^34^

Gene Regulatory Network (GRN) and Transcription Factor (TF) analyses were performed using CellOracle (v 1.18.0)^35^, following procedures outlined in the CellOracle (https://morris-lab.github.io/CellOracle.documentation/tutorials/index.html). Base GRNs were constructed using bulk-ATAC sequencing data from the same cell samples, providing a foundation for inferring regulatory interactions between TFs and their target genes. CellOracle integrated this information with single-cell gene expression data to refine the GRN and identify key TFs driving cellular states and transitions. To further investigate the functional roles of these TFs, knockout (KO) simulations were conducted using CellOracle.

### Statistical analysis

Student’s t-test was performed to evaluate the statistical differences between CM and normal control data sets. For nonparametric data, Mann-Whitney U Test was used. Data was presented as “mean ± S.D.” *p* < 0.05 was considered statistically significant.

## Results

### WTA signatures for CM blood vessels

A total of 123 regions of interest (ROIs), each containing a single blood vessel, were collected for GeoMx WTA analysis. Validation using UEA1 or SMA immunohistochemistry identified seven ROIs containing non-blood vessel structures, which were excluded. An additional 23 ROIs with fewer than 180K aligned reads were removed for quality control, yielding 65 CM vessels (n = 6 patients) and 28 normal vessels (n = 6 subjects) for downstream analysis (Fig. 1A). Morphological parameters assessed per ROI included nuclear counts, average blood vessel wall thickness (BVWT, μm), blood vessel wall area (BVWA, μm²), aligned reads per nucleus, and distance from the epidermis (Supplemental Table 2). Based on these features, vasculature was categorized into five groups (Fig. 1B, C): (1) normal capillaries/venules (17 ROIs), (2) normal small arterioles (11 ROIs), (3) small-thin CM vessels (BVWA < 10,000 μm²; 28 ROIs), (4) large-thin CM vessels (BVWT < 70 μm, 1–2 cell layers, BVWA > 10,000 μm²; 11 ROIs), and (5) large-thick CM vessels (BVWT > 70 μm, >3–4 cell layers, BVWA > 10,000 μm²; 26 ROIs).

**Figure 1.**
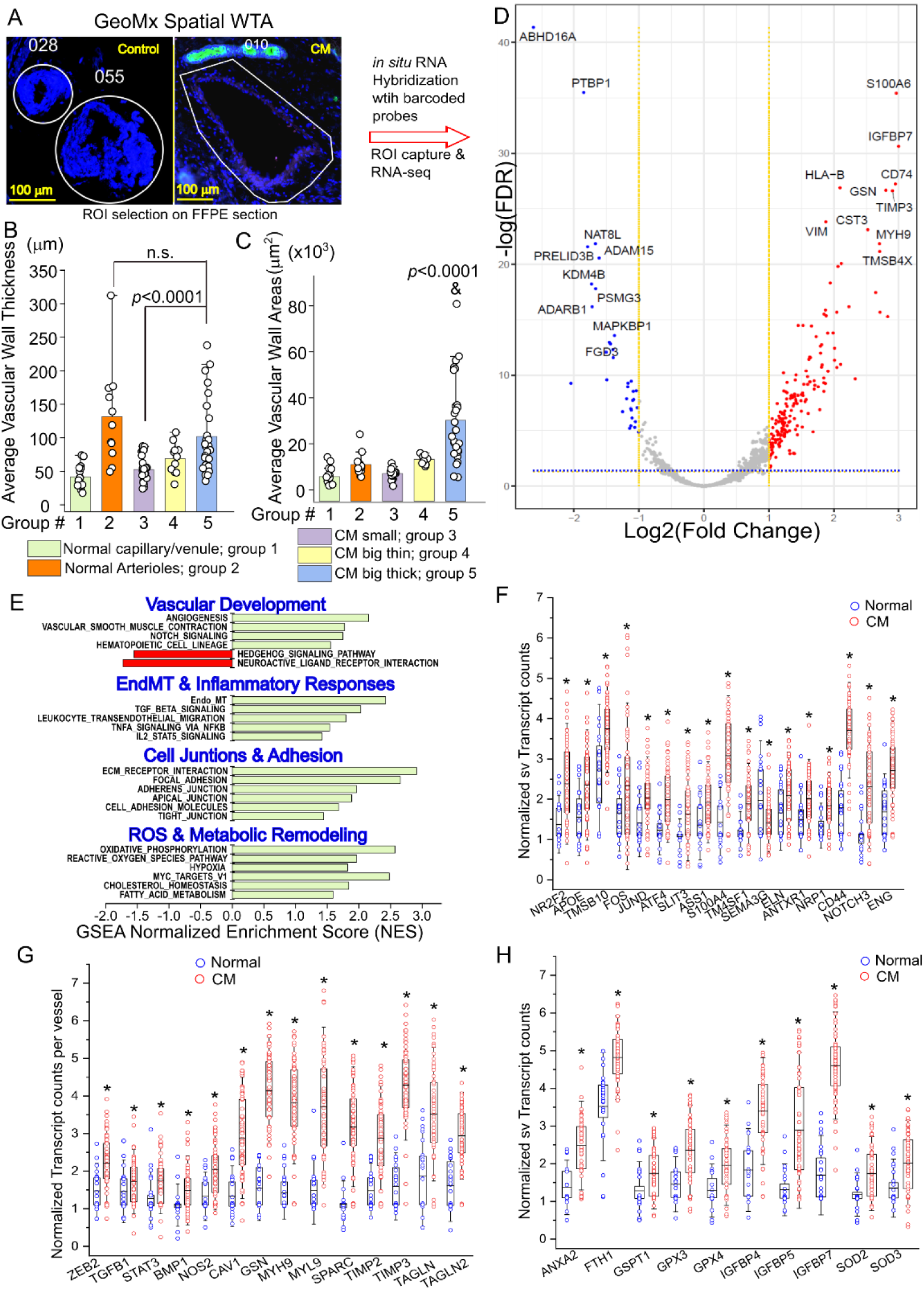
WTA profiles for CM vs normal vasculature: A, Each ROI image contains a single blood vessel from FFPE sections in normal control skin or CM lesion for WTA profiling using a GeoMx platform; B-C, Morphological features of blood vessels in ROIs; The profiled vasculatures were categorized into five groups based on their average vascular wall thickness and wall areas; D, Volcano plot for DEGs from WTA profiles of CM vs normal control blood vessels; Red or blue dots in volcano plots: significantly up- or down-regulated DEGs, respectively (FDR < 0.05). E, Single vessel normalized enrichment score for GSEA (svNES_GSEA) showing the major dysregulated pathways in CM vs normal dermal vasculature; F-H, Representative DEG plots related to EC and EPC biomarkers (F), EndMT panel (G), and Hypoxia/ROS signaling (H). Each dot represents a single vessel. * FDR < 0.05.

There were 467 upregulated and 136 downregulated DEGs in CM as compared to normal vasculatures (Fig 1D, Supplemental Fig 1). Single vessel (e.g., each ROI) Gene Set Enrichment Analysis (svGSEA) showed dysregulated pathways mainly involved vascular development (angiogenesis, Notch signaling, Hematopoietic cell lineage, and hedgehog signaling) and EndMT related biological processes (inflammatory responses, cell junctions and adhesion, and hypoxia/ROS signaling) (Fig 1E, Supplemental Fig 1).

### Endothelial NR2F2 coincides with CM pathological progression

CM ECs displayed features of EPCs,^12,14^ co-expression of both venous and arterial biomarkers, and loss of arteriole-like vasculature, indicating impaired arterial differentiation.^12^ The WTA data identified many DEGs related to venous (NR2F2, TMSB10, JUND), arterial (ASS1, S100A4, TM4SF1, ELN), and progenitor (CD44, NOTCH3, KLF2, ENG) biomarkers (Fig 1F), suggesting disrupted vascular lineage determination in CM.

The IF validation of cellular colocalizations and patterns of these DEGs related to EC and EPC biomarkers were shown Supplemental Figs 2-3. Of note, NR2F2 is a TF for determining venous identity,^36^ and ASS1 is a post-capillary arterial biomarker in human dermal vasculature.^37^ We next performed an IF validation crossing CM vasculatures with different sizes in patient biopsies. NR2F2 was found in nuclei of venous ECs while ASS1 was found only in arterial ECs in normal dermal vessels (Fig 2A). Scattered SMCs showed a strong nuclei NR2F2 expression in perivascular regions in both venules and arterioles (Fig 2, Supplemental Figs 4 and 5). In lesional vasculatures with small sizes, many ECs co-expressed NR2F2 and ASS1 (Fig 2B, Supplemental Fig 4). In intermediate-sized lesional vessels, there were several expression patterns in ECs within single vessels: NR2F2 positive, ASS1 positive, and weak positive or negative for both biomarkers (Fig 2B, Supplemental Fig 4), demonstrating a dynamic heterogeneity of (1) presence of both venules and arteriole-like ECs within one vessel, and (2) co-existing of both venous and arterial signatures in the same ECs. However, in thick, large-thick vessels with multiple layers of SMCs, only NR2F2 was observed in the nuclei of many ECs but not ASS1 (Fig 2B, Supplemental Fig 4). Neither NR2F2 nor ASS1 was observed in ECs in large-thin vessels with one or two layers of SMCs. The proportion of ECs with positive NR2F2 nuclei per vessel in CM was significantly higher than normal dermal blood vessels (Fig 2C). CM ECs express stem cell marker CD133.^12^ CD133 was localized in plasma membranes and NR2F2 was in nuclei in CM ECs (Supplemental Fig 5), consistent with progenitor-like states. The dynamics and transition patterns of endothelial NR2F2 were outlined in Fig 2D, coinciding with progressive enlargement of lesional vasculature.

**Figure 2.**
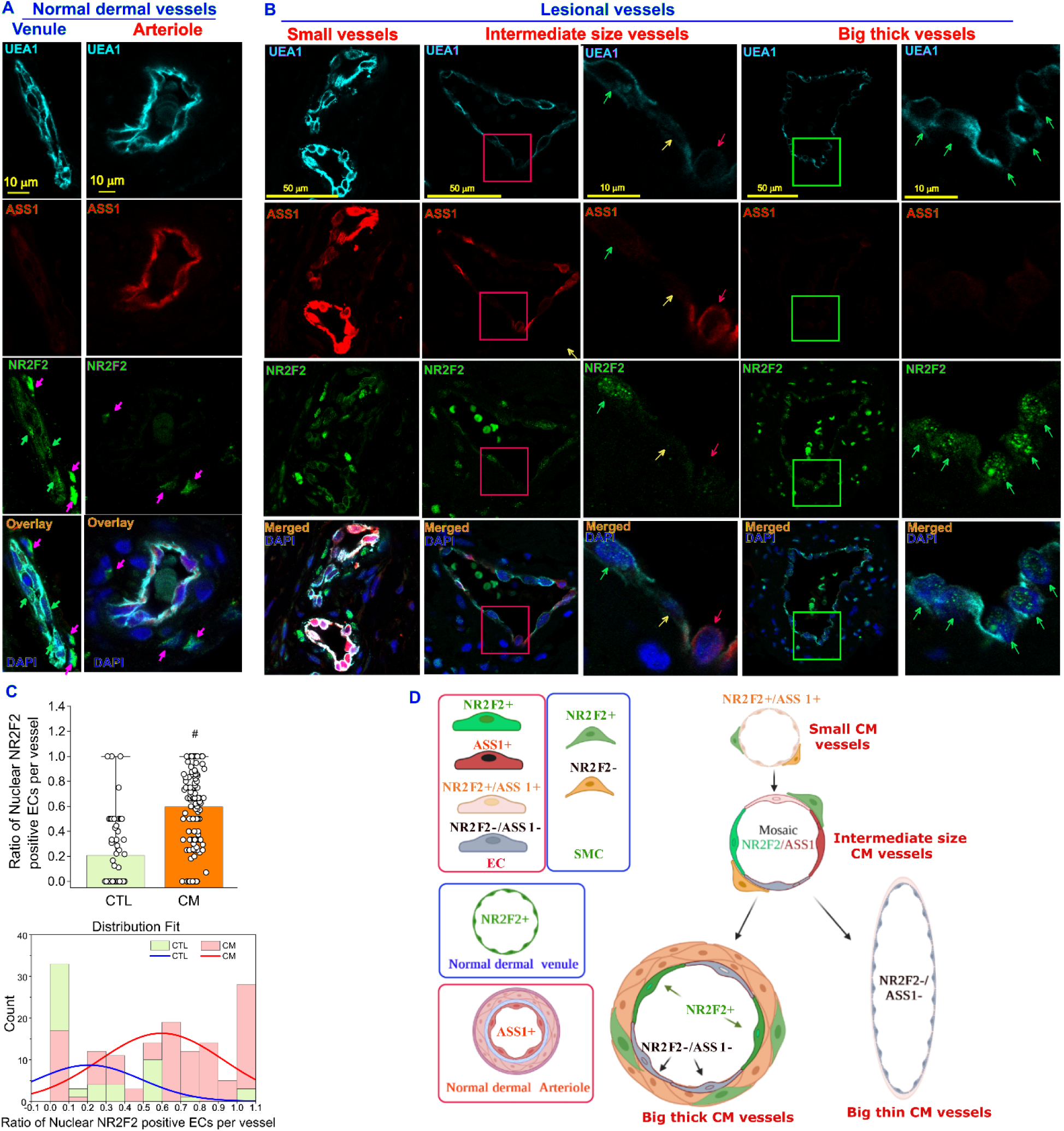
Endothelial NR2F2 coincides with pathological progressions of CM vasculature. A, venous driver NR2F2 (green) was found in nucleus of ECs in normal dermal venous, but dermal arterial biomarker ASS1 (red) was mainly present in arterial ECs in normal dermis. Some scattered SMCs showed strong nucleus NR2F2 IF signaling in both venous and arterioles (pink arrows). B, In small size CM vessels, the co-expression of ASS1 and NR2F2 were observed in many ECs. In intermediate size vessels, ECs in one vessel could show heterogenous patterns of NR2F2 (green arrows) positive, ASS1 (red arrows) positive, or negative for both biomarkers (yellow arrows). In big dilated and thick lesional vessels, NR2F2 but not ASS1 was found in the nucleus of many ECs. The 2^nd^ panel of intermediate size or big size vessels is a high magnification of the box area of each image in the 1^st^ panel, respectively. UEA1 (cyan) staining was used to show ECs. C, Quantitative analysis and distribution fit of the ratio of nuclear NR2F2 positive ECs per vessel in normal skin as compared to CM. #, *p* < 0.0001. D, the dynamics and transition patterns of nuclear NR2F2 in ECs coincides with progressive enlargement of lesional vasculature.

svGSEA highlighted enrichment of lipid metabolism pathways (Fig. 1E). IF staining validated altered expression of APOE and FADS2, two regulators of endothelial lipid metabolism. APOE involves in endothelial dysfunction and FADS2 is expressed in dermal ECs engaging cholesterol synthesis.^38,39^ In normal skin, dermal capillaries or venules expressed low or moderate levels of FDAS2 but not APOE; normal dermal arterioles expressed high levels of both APOE and FDAS2 (APOE^H^/FDAS^H^, Supplemental Fig 6). In CM lesions, three types of vessels were observed, low levels of APOE and high levels of FDAS (APOE^L^/FDAS^H^), high levels of APOE and low levels of FDAS APOE^H^/FDAS^L^, and APOE^H^/FDAS^H^ (Supplemental Fig 6), suggesting heterogeneous metabolic reprogramming linked to impaired vascular lineage specification.

### Molecular signatures for enlarged CM blood vessels

Principal Component Analysis (PCA) of GeoMX samples revealed distinct clustering between normal and CM vessels (Fig 3A). During development, both dermal arterioles and venules are differentiated from primitive capillary plexus (PCP).^40,41^ In default, PCP is thought to develop into a venule; while turning off venule specific genes and switching on arteriole specific genes will drive dermal PCP differentiation into arterioles.^40^ The normal arteriole samples were a part of cluster within capillary/venule samples, reflecting normal PCP-to-arteriole differentiation. The samples of CM big-thin (group 4) and big-thick (group 5) formed two separate clusters within the small CM vessels (group 3) (Fig 3A).

**Figure 3:**
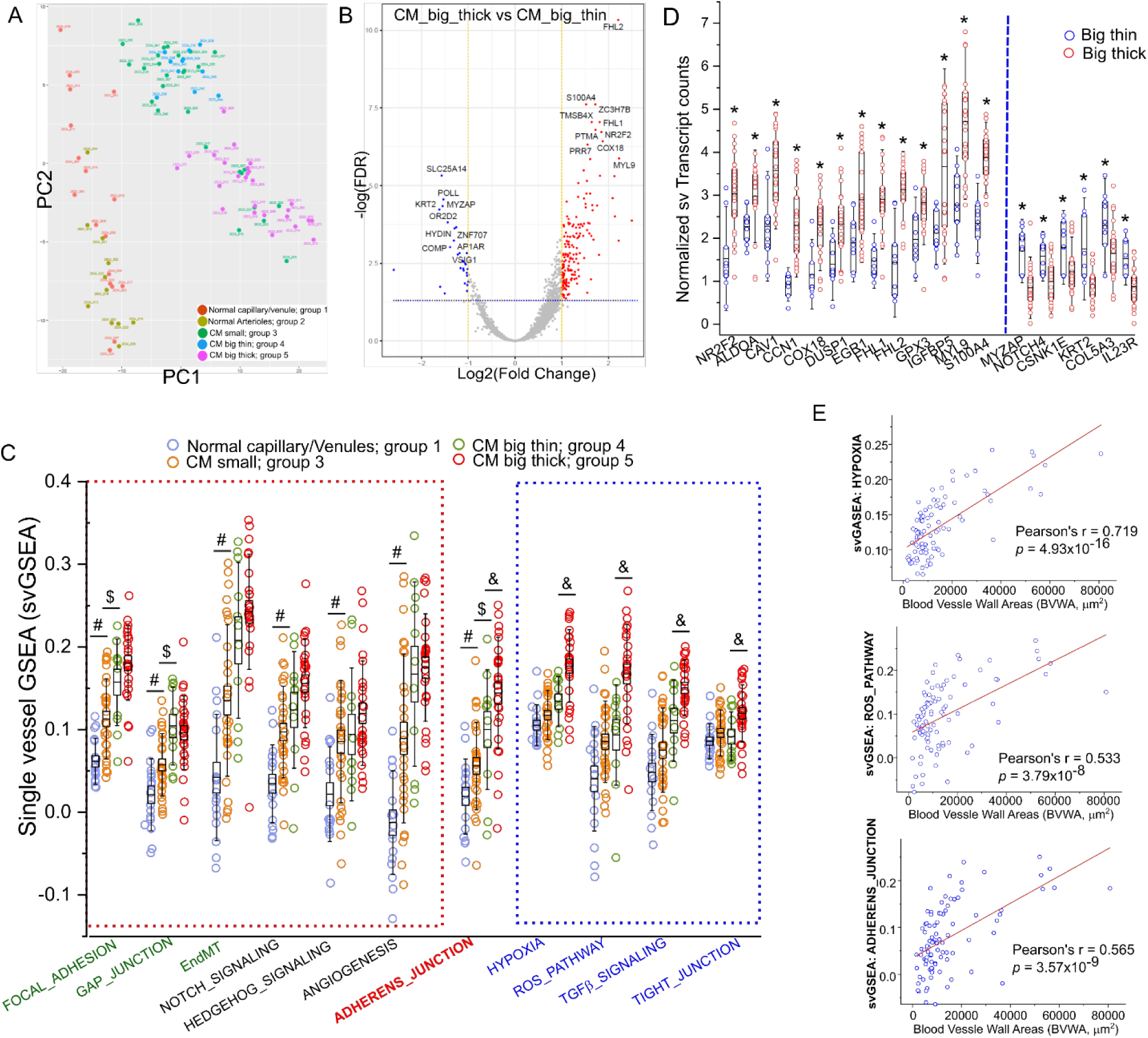
WTA profile comparison of big thick versus big thin CM vessels. A, PCA plot showing sample distances among all profiled blood vessels in ROIs from groups 1-5. B, Volcano plot for DEGs from WTA profiles of big thick versus big thin CM vessels; Red or blue dots in volcano plots: significantly up- or down-regulated DEGs, respectively (FDR < 0.05); C, Scattered plot showing svNES_GSEA for main pathways among groups; # group 3 vs group 1 with FDR < 0.05; $ group 4 vs group 3 with FDR < 0.05; & group 5 vs group 4 with FDR < 0.05; D, Representative DEGs of WTA profile comparison of big thick versus big thin CM vessels. * FDR < 0.05. E, Pearson correlation between BVWA and svNES_GSEA of hypoxia, ROS, and AJ pathways, respectively. Each dot in panels C-E represents DEG or GSEA from a single vessel.

We then performed subgroup comparisons between CMs (groups 3-5) and normal ones (groups 1-2) (Supplemental Figs 7-13). There were 59 downregulated and 362 upregulated DEGs in big-thick (group 5) versus big-thin CM vessels (group 4), including NR2F2 and many vascular remodeling biomarkers (Fig 3B, Supplemental Fig 14). The svGSEA demonstrated that junction pathways, EndMT, Notch, Hedgehog, and angiogenesis were significantly enriched in small CM vessels (group 3) versus normal dermal capillaries/venules (group 1) (Fig 3C), suggesting their functional involvements of differentiation. Inflammatory pathways (hypoxia, ROS, and TGFβ) and tight junctions (TJs) were significantly enriched in big-thick versus big-thin CM vessels (Fig 3C), implicating their functions in EC remodeling and disease progression. Interestingly, AJs were significantly dysregulated across both comparisons (Fig 3C). Some presentative DEGs for big- thick or big-thin CM vessels were shown in Fig 3D. IF validation showed that MYOZAP was observed in both ECs and SMCs. It exhibited fragmented and distorted immunostaining patterns in big-thick versus big-thin vessels (Supplemental Fig 15). IL23A1 expression was mainly observed in SMCs but not ECs with similar immunostaining signal intensities in both types of CM vessels (Supplemental Fig 15). We next used the blood vessel wall area (BVWA) as the morphological index of blood vessel thickness and size. The correlation analysis showed that BVWA was significantly correlated with scGSEA scores of hypoxia, ROS, and AJs (Fig 3F), suggesting their roles in the pathological progression of CM blood vessels with thick versus thin walls.

### Junctional impairment as an EndMT phenotype in CM

SvGSEA showed EndMT was enriched in the CM vasculature compared with normal dermal blood vessels (Fig 1E). One landmark phenotype of EndMT is dysregulations of EC-EC junctions including TJs and AJs. We performed re-analysis of our existing TEM data of CM vessels.^13,42^ CM ECs showed a wide variety of impairments of TJs and AJs, including fragmented and tortuous morphology, reduction or breakdown of electron density, widened AJ spacing, and absence of entire junctional structures (Fig 4A, B). The overall ratios of impaired endothelial TJs and AJs in CM lesions (83.99% ± 24.66%, n= biopsies from 12 patients) were significantly higher than the baseline in normal skins (4.66% ± 8.77%, n= biopsies from 12 subjects, *p*<0.0001) (Fig 4C). Half of the AJs (49.9%) in CM vasculatures versus 4.1% AJs in normal dermal vessels, showed junctional spacing larger than 20 nm between adjacent ECs (Fig 4D, E; *p*<0.0001).

**Figure 4:**
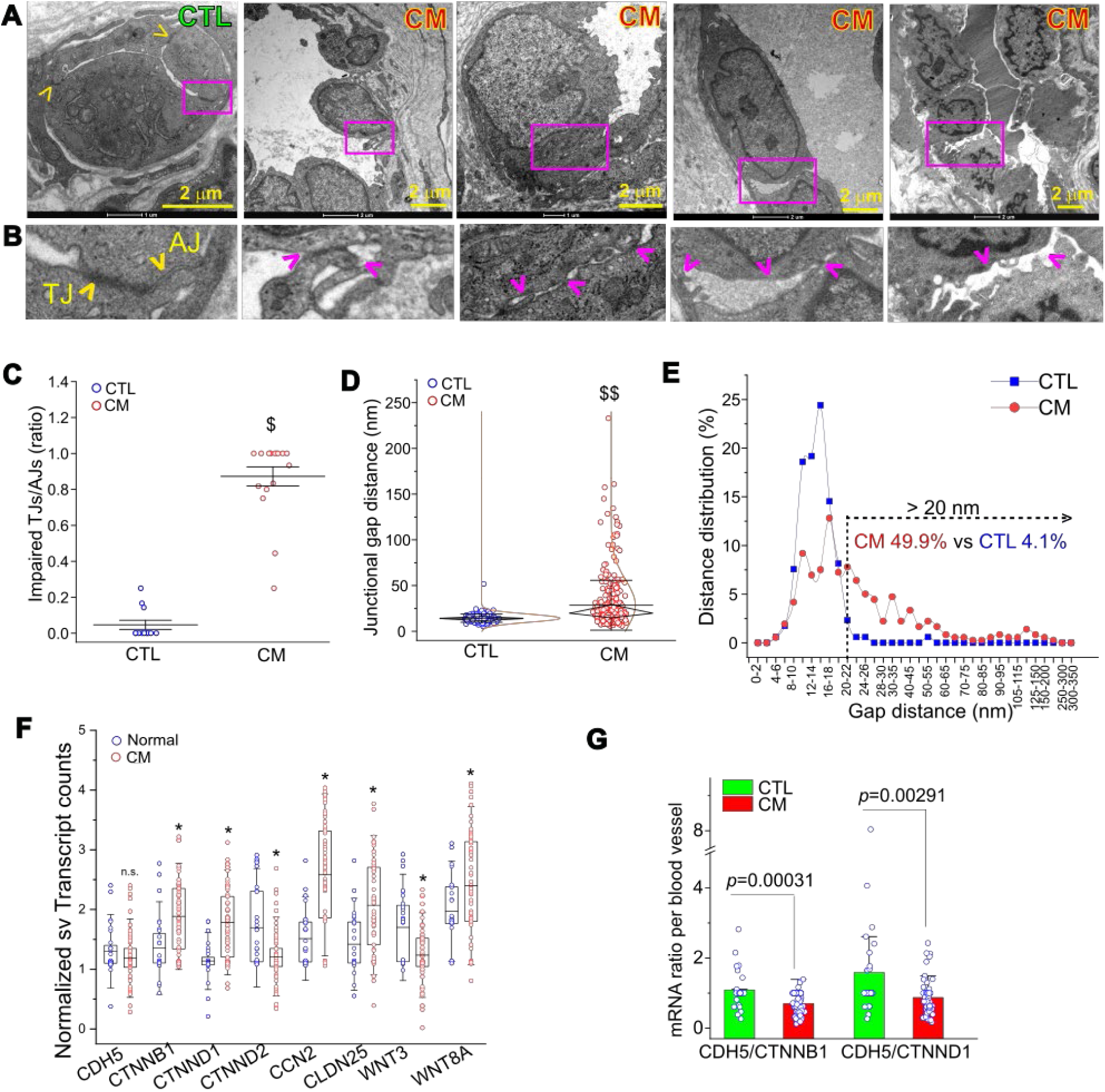
Impairments of AJs in CM vasculature. A, TEM showing the AJs among ECs from a normal dermal capillary and various types of fragmented and tortuous AJs among CM ECs with enlarged junctional spaces. B, The magnified box area from upper panel in (A) respectively. Yellow arrowhead: normal TJ and AJ; Pink arrowhead: impaired AJs. C, The ratio of impaired TJs and AJs over normal ones among control or CM vessels. Each dot represents one subject or patient. $ p<0.0001. D, Junctional gap distances in AJs from normal and CM vessels. $$ p<0.0001. E, The distribution of gap distances in AJs from normal vessels as compared to CM vessels. F, Scattered plot showing the normalized transcript counts of AJ-related genes in CM as compared to normal vessels. Data was from GeoMx WTA profile. n.s., no significance for CDH5; * FDR < 0.05. G, mRNA ratio of CDH5/CTNNB1 or CDH5/CTNND1 per blood vessel in control as compared to CM lesions.

To further characterize the impairments of AJs in CM vasculatures, we examined the transcriptime atlas of AJ-related factors including CDH5, CTNND1, CTNNB1. Both CTNND1 and CTNNB1, but not CDH5, were significantly upregulated in CM blood vessels than normal vessels (Fig 4F). The ratios of CDH5 to CTNND1 or CTNNB1 per vessel were significantly lower in CM vessels than in normal vessels (Fig 4G). IF staining showed the co-locolization of both CTNND1 and CDH5 among EC-EC contacts (Supplemental Fig 16), further valiation was performed by immunoblotting using cell lysate from the whole biopsy skins (Supplemental Fig 17). These data demonstrate molecular bases underlying AJ impairments in CM vasculature.

We next validated the CTNND1 patterns in lesional blood vessels using IF staining. Normal dermal capillary and venous ECs are flattened with orientations along the vessel’s long axis with intact cell-cell junctions (Figs 5A, B). CM ECs exhibit various morphological phenotypes: (1) flattened shape ECs orient along the vessel’s long axis, and (2) cuboidal shape ECs orient perpendicularly along the vessel’s long axis with frequently impaired EC-EC contacts (Figs 5C-H). Both types of ECs can be found in CM vasculature of all sizes. Normal human dermal capillary ECs had either barely detectable CTNND1 IF signals [Type 1 or normal 1 (N1)] (Fig 5A, Supplemental Fig 18A) or puncta CTNND1 (CTNND1^pn^) [Type 2 or normal 2 (N2)] among normal venous EC AJs (Fig 5B; Supplemental Fig 18B). Occasionally, a few CTNND1^pn^ ECs were found in the lesions (Supplemental Fig 18C; 2). In general, CM ECs in flattened shapes had CTNND1 IF signals covering the basal membranes (CTNND1^bm^) (Type 3; Fig 5C; 3; Supplemental Fig 18); while CM ECs in cuboidal shapes usually showed CTNND1 IF signals on the lateral membranes (CTNND1^lm^) (Type 4; Fig 5C; 4; Supplemental Fig 18) or the lateral basal membrane (CTNND1^lb^) (Type 5; Fig 5G; 5; Supplemental Fig 18). Some ECs with a full transition of columnar shape exhibited an extended lateral membrane CTNND1 pattern (CTNND1^elm^) (Type 6; Fig 5E; 6; Supplemental Fig 18). These EC morphological features suggest that CTNND1^elm^ is a pathologically advanced varaint of CTNND1^lm^. A few ECs developed a pattern of CTNND1 in all membranes (CTNND1^all^) (Type 7; Fig 5G; 7; Supplemental Fig 18). We did not find any obvious CTNND1 IF signals in SMCs. This data suggests that heterogenous CTNND1 patterns present a snapshot of the dynamics of CM EC remodeling during disease progression.

**Figure 5:**
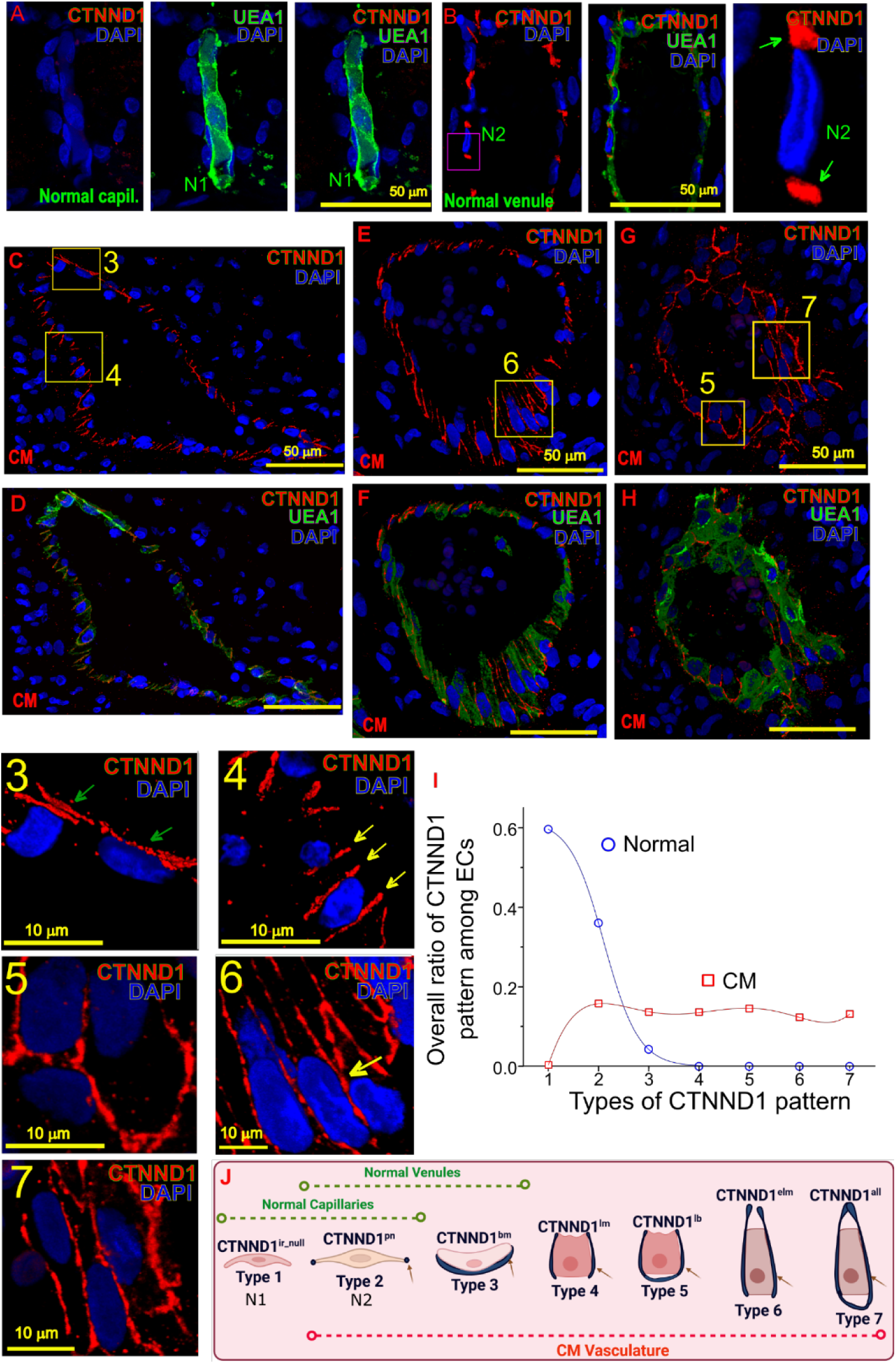
CM ECs with heterogenous CTNND1 patterns reflecting dynamics of EC phenotype remodeling. A and B, Normal human dermal capillary ECs had either barely detectable CTNND1 (red) IF signals (A, normal type 1, N1) or puncta CTNND1 (CTNND1^pn^) (B, normal type 2, N2) between normal venous ECs. The right panel showing a high magnification of the red box area of N2; C-H, CM ECs represent various patterns of CTNND1 among lesional blood vessels. Numbered boxed areas in (C), (E), and (G) are magnified and shown in the individual panels with the same number, presenting heterogenous types of CTNND1 patterns. UEA1 (green) staining was used to show the morphologies of vasculature in (D, F, and H). Z-stacks confocal images were processed for CTNND1 subcellular patterns. I, Overall ratios of various CTNND1 patterns among ECs in normal dermal and CM blood vessels. J, Predicted trajectory of endothelial CTNND1 patterns that reflects the diversity and dynamics of CM EC remodeling during disease progression.

### Vascular identity transition as an EndMT phenotype in CM

The second landmark phenotype of EndMT is the cellular transdifferentiation in which ECs partially lose their identity and acquire mesenchymal cell-like features, including a spindle-shaped, elongated morphology (such as remodeled ECs with CTNND1 patterns of types 4 to 7) or/and the acquisition of contractile properties. The representative EndMT related DEGs, including ZEB2, TIMPs, IGFBPs, BMP1, MYLs, and TAGLN (a.k.a., SM22) were shown in Fig 1G. We observed active intermediary EndMT zones within lesional vessels (Fig 6A, B). Such an EndMT zone was formed by several ECs exhibiting cuboidal morphology with CTNND1^lm^ (type 4) and CTNND1^elm^ (type 6) patterns. They presented some SMC phenotypes, e.g., expression of aSMA (Fig 6C, supplemental Fig 19), suggesting the acquisition of myogenic tone. These remodeled ECs present both EC and SMC biomarkers, demonstrating an active and intermediary EndMT remodeling phase in lesional blood vessels.

**Figure 6:**
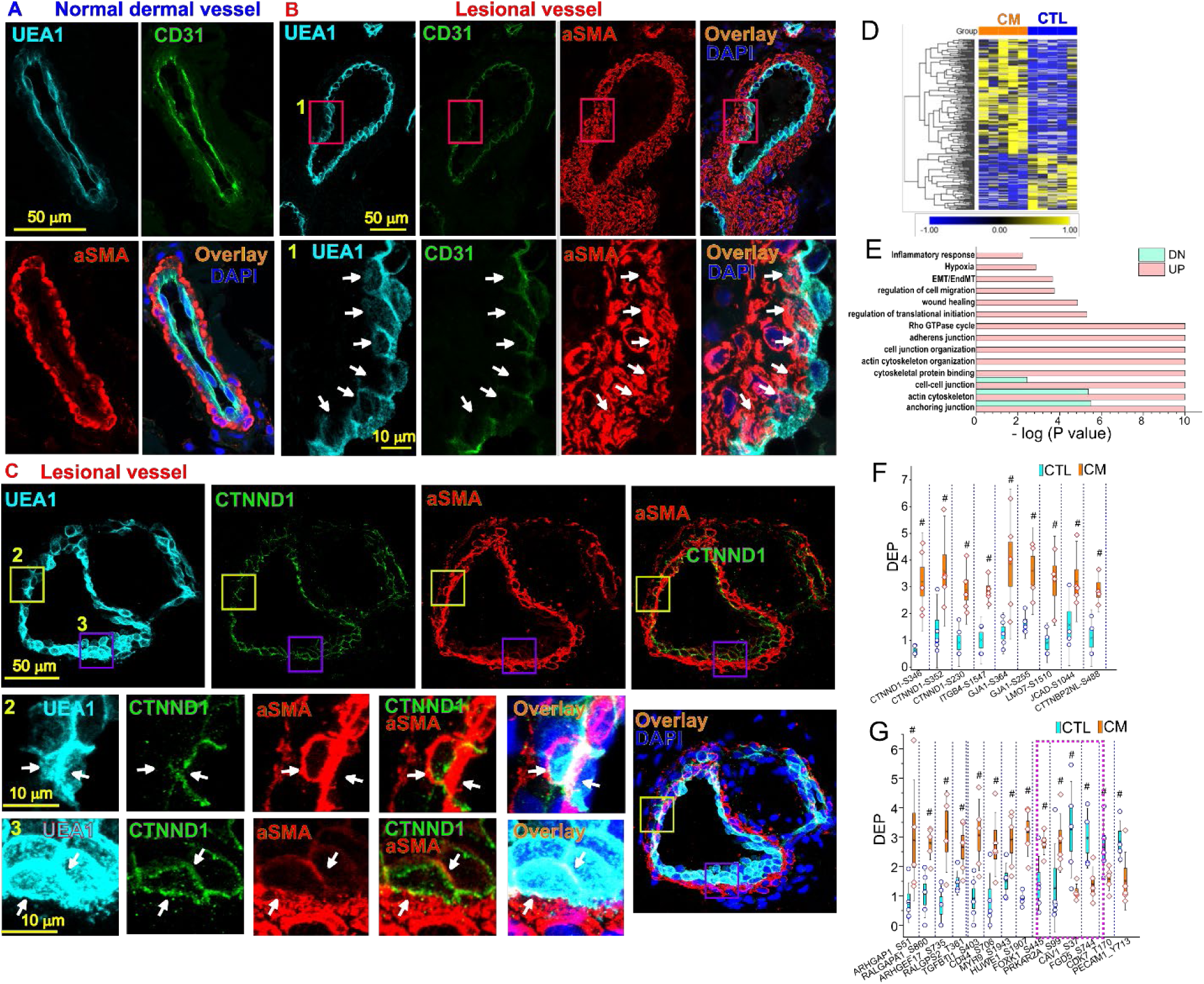
CM ECs undergoing EndMT hallmarks and phosphoproteomics profiles in CM lesions. A, A normal human dermal vessel exhibits distinct aSMA (red) signals in juxtaposition to ECs showing by UEA1 (cyan) or CD31 (green); B, A CM lesional vessel develops an EndMT zone (red box 1) containing remodeled ECs (white arrows). Those ECs have acquired phenotypes of SMCs, e.g., expressing aSMA and showing cuboidal shapes. In (B), lower panels are a high magnification of the red box areas (box 1) in upper panels. DAPI: blue. C, ECs in active EndMT zones presenting various endothelial CTNND1 patterns in a CM vessel. Lower small panels 2 and 3 are a high magnification of the boxed areas with the same number in the upper panel showing endothelial CTNND1^lb^ (yellow box 2) and CTNND1^all^ (purple box 3, multiple layers of ECs) patterns. White arrowhead: the colocation of UEA1 (cyan), CTNND1 (green), and SMA (red). D, Heatmap data showing representative DEPs in CM lesions as compared to normal skin; E, Significantly enriched pathways involving DEPs’ functions; F, Scattered plot showing DEPs of junctional related proteins including CTNND1 at S346, S352, and S230 sites in CM lesions vs normal controls (n=5 subjects); G, Scattered plot for some representative DEPs related Rho GTPases and endothelial functions.

Metabolic remodeling is another EndMT landmark phenotype. We next validated several DEGs related to EndMT and metabolic remodeling using IF staining in CM vasculature, including VIM, MYH9, TWIST1, TM4SF1, SERPINE1, TIMP3, Annexin V, eNOS, LRP1, SPARC, Ki67, GCLM, S100A4, and HIF1α (Supplemental Figs. 5, 20, 21 and 22). In addition, the transcripts of Collagens 1, 3, 4, 5, 6, and 14, another major category of EndMT biomarkers, were systemically and significantly upregulated in CM vessels compared with normal dermal vessels (Supplemental Fig 23A).

### Phosphorylated modulations of junctional proteins in CM

Next, we screened the global activation of cellular signalings in CM lesions by mass spectrometry (MS) to reveal phosphoproteomics and identify differentially expressed phosphorylated peptides (DEP) (n=5 normal and 5 patients). A total of 2,044 phosphorylated proteins (including 4,779 phosphorylated sites) were identified with high confidence (localization probability > 0.75). The relative quantitation of DEPs was divided into two categories: up-regulation (fold-change (FC) > 1.5 and p < 0.05) and down-regulation (FC < 0.667 (1/1.5) and p < 0.05). There were 203 upregulated DEPs (158 proteins) and 100 downregulated DEPs (88 proteins) in CM lesions compared with the normal control skins (Fig 6D; Supplemental table 3). Approximately 51 out of 158 upregulated phosphorylated proteins had biological functions entangled to EMT/EndMT and its related cell-cell junction, Rho GTPase cycle, and anchoring junction (Fig 6E). Other enriched functional groups of DEPs included inflammation, cell migration, and cytoskeleton (Fig 6E), which were also interactive with EndMT. The representative phosphorylated cell junction proteins in the lesions were CTNND1, TJP1, GJA1, and JCAD (Fig 6F). In particular, CTNND1 had DEPs at S230, S352, and S346, exhibiting 2.3, 3.7, and 8.9-fold increases compared to the control skin (*p*=0.03, 0.01, and 0.002), respectively (Fig 6F). The profound DEP signatures also enriched in GTPases, for example, ARHGAP32 (S706), ARHGAP1(S51), RALGAPA1(S860), ARHGEF17 (S735), and RALGPS2 (T361) in CM lesions than in normal skin (Fig 6G). Functional analysis of DEPs showed the convergence of pathways in cell-cell junctions, anchoring junctions, Rho GTPases, EMT/EndMT, hypoxia signaling, and inflammatory responses (Fig 6E, Supplemental Fig 23B), consistantly with GeoMx WTA data.

### Impaired trajectory of CM iPSC differentiation into iECs

We next performed scRNA-seq analysis to gain mechanistic insight into the CM pathological progression. Samples were collected on days 0, 4, 8, and 15 during differentiation course from iPSCs to iECs (Fig 7A). Ten major distinct populations, e.g., iPSC, iPSC_MC, MC_1, MC_2, MC_3, SMPC, MC_EPC, SMC_1 (contractile phenotype), SMC_2 (synthetic phenotype), and EC were clustered (Figs 7B-C, Supplemental Fig 24A). The associated biomarkers of each cluster were plotted in Fig 7D. We next compared the differentiation trajectories of normal and CM samples using pseudotime analysis. Both normal and CM iPSCs showed two distinct differentiation pathways, e.g., endothelial lineage and SMC lineage (Fig 7E). The differentiation of endothelial lineage was observed in a pathway from clusters of iPSC_MC, MC_1, EPC, to EC (Figs 7F, G). While the differentiation of SMC lineage followed the track from clusters of iPSC_MC, MC_SMPC, SMC_2, to SMC_1 (Figs 7 H, I). CM iPSC yielded a lower percentage of iECs (25.9% versus 39.6%) but resulted in a higher ratio of iSMC lineages (SMC_1 plus SMC_2: 14.1% versus 8.6%) than normal ones (Figs 7F-I). Our iEC differentiation protocol is in favor of arterial lineage commitment, demonstrating by high level expression of arterial biomarkers in iECs including CXCR4, Hey1, Hey2, NOTCH1, DLL4, and EFNB2 (Fig 7D, Supplemental Fig 24B). These data suggest that CM iPSCs have a reduction in arterial EC lineage differentiation but an enhancement in SMC lineage determination (Fig 7J), which is consistent with our previous report showing impairments of arterial differentiation in CM vasculature.^12^

**Figure 7:**
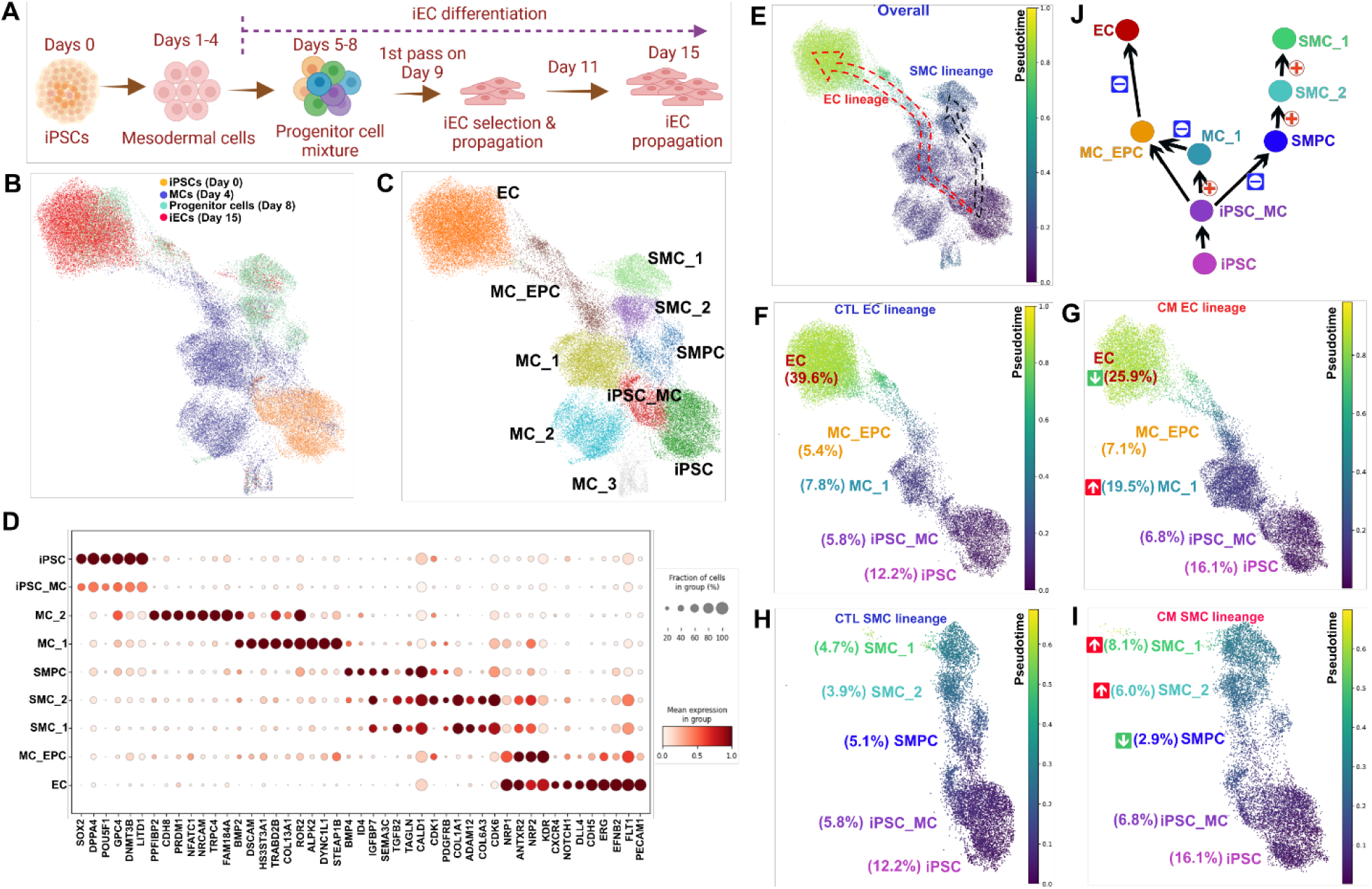
Differentiation impairments of CM iPSC to vascular lineages. A, Schematic of normal or CM iPSCs differentiation to iECs. Samples were collected on days 0, 4, 8, and 15 for scRNA-seq analysis. B and C, UMAP showing the distributions of cell cultures among sampling times and cell types. D, Dot plot showing biomarkers for different clusters of cell types. E, Pseudotime analysis showing differential trajectories of EC and SMC lineages. F and G, Pseudotime analysis showing a differential stall of EC lineage in CM as compared to normal iPSCs with an increased MC_1 subpopulation but increased EC subset. H and I, Pseudotime analysis showing a differential enhancement of SMC lineage in CM as compared to normal iPSCs with increased SMC_2 and SMC_1 subsets. J, Schematic of differential trajectories and impaired paths in CMs.

To identify potential TFs underlying these differential impairments of CM iPSCs, we constructed base GRNs of normal and CM iECs from the bulk ATAC-seq data (Fig 8A). The degree centrality calculated using CellOracle (v 1.18.0) was used to rank the top TFs through GRNs in normal and CM clusters of MC_1, EPC, EC, SMPC, SMC_2, to SMC_1 (Figs 8B, C; Supplemental Figs 24C). To explore the role of TF activity on cell-specific gene expression profiles, we performed *in silico* KO analysis using CellOracle. In each perturbative simulation, the changes of vector field, a metric indicating the shift of cell identity, and perturbation score (PS), a metric indicating the strength of shift, were used to predict the KO effect of individual TFs. A systemic KO simulation identified top TFs with greater or less impacts on cell identity shifts in each cluster in CM and control (Figs 8B, C; Supplemental Figs 24C). There were a handful of TFs that overlapped with the DEGs from GeoMx data, including KLF2, NR2F2, EPAS1, NR4A1, ID1, ID4, FOS, HEY2, TAL1, JUN, MSX2, PAX8, NPAS3, MEIS3, and RFX2 (Figs 8D-I; Supplemental Figs 25 and 26). For example, KLF2 KO suppressed MC_EPC cluster formation but promoted differentiation from MC_EPC to EC cluster in CM compared with the control (Figs 8D, H). NR2F2 and EPAS1 KOs exhibited inhibition on SMPC cluster in both CM and control (Figs 8E, F, H). NR4A1 KO inhibited SMPC cluster in the control but promoted SMPC cluster in the CM (Figs 8G, H). KLF2, NR2F2, and EPAS1 were upregulated, whereas NR4A1 was downregulated in GeoMx profile (Fig 1F, Fig 8I). Together, these data suggest that upregulation of NR2F2 and EPAS1, and downregulation of NR4A1 may drive the SMC phenotype development in CMs. The KO simulations of other overlapping TFs in GeoMx DEGs were performed. In summary, the following TFs were predicted to have more engagements in EC lineage development in CM than control: ID4, FOS, HEY2, JUN, TAL1, and NPAS3. While the TFs including MSX2, RFX2, MEIS3, NPAS3, and PAX8 contributed more to SMC phenotype development in CM than in the control (Supplemental Figs 25 and 26).

**Fig 8:**
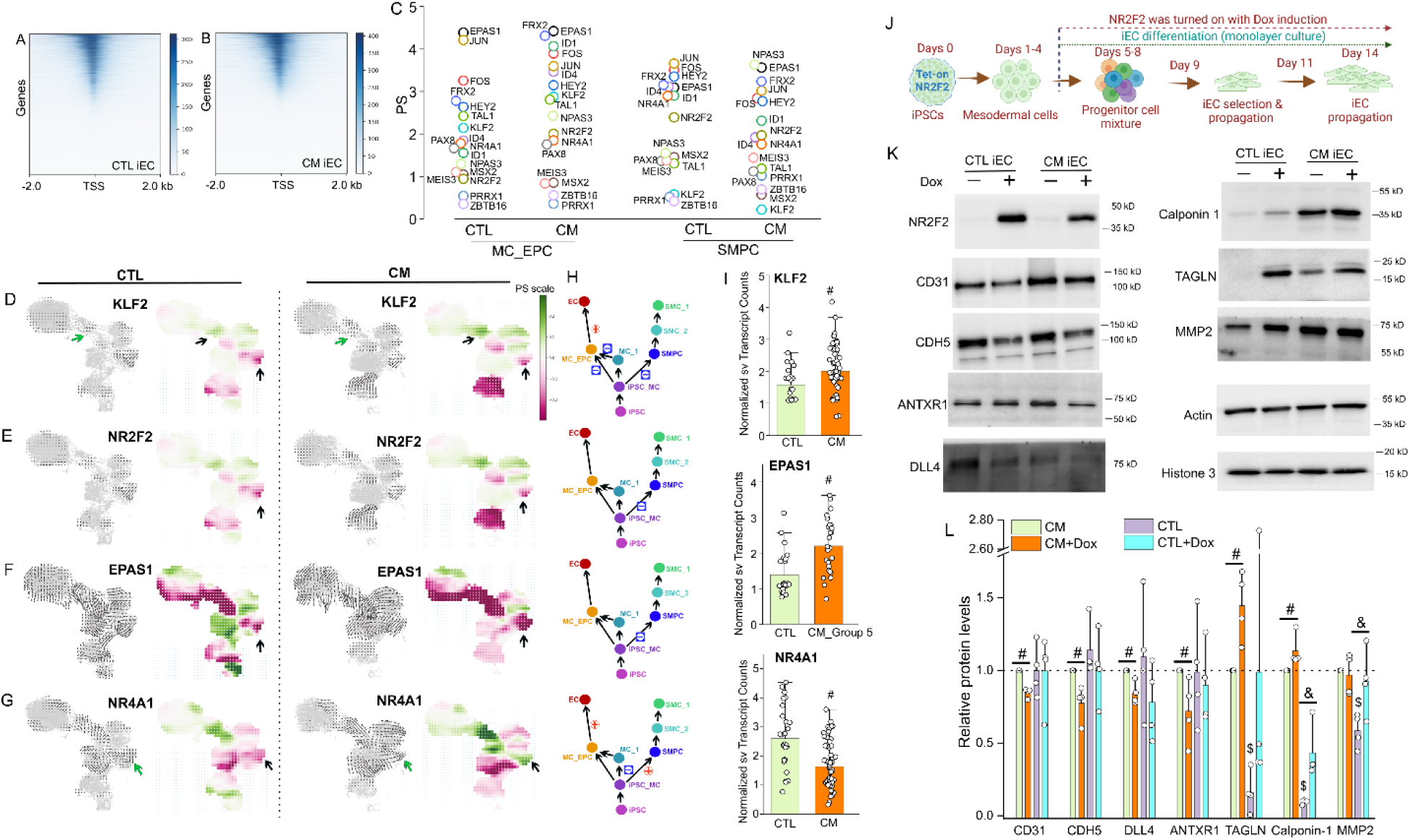
In silico gene perturbation and forced expression of NR2F2 promoting EndMT. A and B, Heatmaps showing enrichment at transcription start site (TSS) for the ATAC-seq profiles of normal (A) and CM iECs (B). GRN models were constructed from peaks files of ATAC-seq data. C, perturbation scores of representative TFs among GeoMx DEGs showing their impacts on cell identity shifts in clusters of MC_EPC and SMPC in CM and CTL through KO simulation. D-G, CellOracle simulation of cell-state transition in *Klf2* (D), *Nr2f2* (E), *Epas1* (F), and *Nr4a1* (G) KO simulation in CTL and CM. Summarized simulation vector field and the perturbation scores were shown, respectively. PS scale, green color indicates differentiation promoted; red color, differentiation suppressed with specific TF KO simulation. H, Schematic of impact on perturbative shifts for iEC or iSMC lineages with each specific TF KO simulation in CM and CTL. I, Single vessel GeoMx profile of specific TFs in CM as compared to CTL. #, FDR < 0.05. J, Schematic of normal or CM iPSCs with integration of a Tet-on system to induce over-expression of NR2F2 with Doxycycline (Dox). Dox was added on day 5 to day 14 during iEC differentiation stages. K, Forced expression of NR2F2 at the stages from EPCs to iECs could cause reductions of EC biomarkers such as CD31, CDH5, DLL4, ANTXR1, and increases in EndMT biomarkers such as TAGLN, Calponin 1, and MMP2 in iECs. L, Relative protein levels of selective EndMT biomarkers. The basal level of each protein in CM iECs is set as 1 (dashed line). #, *p*<0.05 Dox versus mock in CM iECs; &, *p*<0.05 Dox versus mock in CTL iECs; $, *p*<0.05 basal level in CM versus CTL iECs. N=4 independent experiments. The average of normalizations to two housekeeping proteins β-actin and histone 3 is set as relative protein level.

### NR2F2 promotes iECs acquiring EndMT-like SMC phenotypes

NR2F2 is a master TF that determines venous identity and is a major driver for EMT in epdithelial cells. The above NR2F2 KO simulation suggested that NR2F2 promotes SMC phenotype in CM. This led us to explore the role of NR2F2 in the acquisition of EndMT in iECs. We integrated a Tet-on system to induce NR2F2 expression in normal and CM iPSC lines. Dox was added from day 5 (EPCs) to 14 to induce NR2F2 overexpression (Figs 8J, K). CM iECs showed higher basal levels of some EndMT biomarkers such as TAGLN (a.k.a., SM22), Calponin 1, and MMP2 (Figs 8K, L). Forced expression of NR2F2 resulted in decreases in many EC biomarkers, such as CD31, CDH5, ANTXR1, and DLL4 in CM iECs (Figs 8K, L). It also led to increases in SMC biomarkers such as TAGLN and calponin 1 in both CM and CTL iECs. Forced expression of NR2F2 also upregulated MMP2 expressions in CTL iECs, reaching to its’ basal level in CM iECs (Figs 8K, L).

## Discussion

In this study, we have demonstrated that CM vasculature consists of differentially defective ECs. Lesional ECs exhibit several landmark phenotypes of EndMT including AJs impairments, vascular identity transitions and metabolic remodeling. CM iPSCs recapitulate an attenuation of EC lineage differentiation but an enhancement in SMC phenotype development. Consecutive expression of NR2F2 in CM progenitor cells can promote differentially defective ECs to acquire EndMT.

The first evidence showing inflammatory contributors to CMs was reported by Nasim et al.^22^ This study finds that enlarged blood vessels lack mural cells and their perivascular beds contain phagocytic macrophages and neutrophils in brain CMs.^22^ The immune cells such as macrophages have been shown as major contributors to EndMT.^43,44^ In our study, two types of big caliber cutaneous blood vessels are shown in skin lesions: (1) thin ones devoid of SMCs, which are similar to those enlarged ones in the SWS brain,^22^ and (2) thick ones with replications of SMCs, basement membranes, and extracellular matrix such as collagens, as a major vascular type found in CMs. Some ECs in these vessels present predominant EndMT molecular phenotypes upon acquisition of SMC features. Consistently, our WTA data shows a spectrum of inflammatory pathways such as TGF-β, TNFα, ILs, and leukocyte transendothelial migration are upregulated in cutaneous CMs. Soft tissue outgrowth is a clinical manifestation of cutaneous CM lesions associated with EndMT. Our data indicate that lesional ECs likely contribute to fibrosis through the following mechanisms. First, ECs can be direct sources of myofibroblasts through EndMT phenotypical transition. Chronically activated fibroblasts (i.e., myofibroblasts) have been recognized as the main cell population responsible for fibrosis in a variety of disorders.^45^ For example, dermal microvascular ECs from lesional skins in patients with systemic sclerosis present an intermediate stage of EndMT, express both endothelial and myofibroblast markers, and display a spindle-shaped appearance and a contractile phenotype *in vitro*.^45,46^ Second, ECs can secrete various proinflammatory factors to interact with immune cells and release profibrotic mediators to activate fibroblasts. For example, Nasim et al. showed that ECs with the GNAQ (R183Q) mutation can increase the expression of ICAM1, thereby enhancing the adhesion of immune cells on the surface.^22^ An increase in macrophage subset (MRC1/LYVE1) in both brain and skin CMs^22,23^ has suggested the contributive roles of inflammatory components. This study has found that proinflammatory and profibrotic factors such as IGFBP4, 5, and 7 are upregulated in CM vessels, further supporting this mechanism. More interestingly, Nasim et al. has found that there are “sprouting vessels” in CM skin lesions.^23^ We found similar “sprouting vessels” in this study. Morphologically, the tip cells in the “sprouting vessels” exhibited advanced CTNND1 patterns. Therefore, the tip cells in the CM “sprouting vessels” are likely invasive to the SMC layers under an active EndMT process. Collectively, our data suggests that EndMT is detrimental to the pathological progression of CMs.

NR2F2 is a master TF for venous identity.^36^ In addition, the upregulation of NR2F2 increases ROS production^47^ and results in fibrosis^48^ and cardiomyopathy.^47^ NR2F2 has been shown to facilitate the EMT by promoting cell invasion, migration, and expression of CDH2 and vimentin.^49^ However, the role of NR2F2 in EndMT remains unclear. In this study, we demonstrate that NR2F2 is a driver for CM progenitor cells to acquire EndMT: (1) NR2F2 is constantly expressed in a subset of SMCs throughout CM vessels of all sizes; (2) NR2F2 KO simulation shows that NR2F2 can promote SMC phenotype development; (3) Nuclear NR2F2 is consecutively present in CM ECs throughout the disease small caliber vessels to thick large vessel development; and (4) Consecutive expression of NR2F2 can lead to EndMT acquisition in CM iECs. Our data suggest that hypoxia and ROS signaling are the driving pathways for EndMT in CMs. The overexpression of NR2F2 can increase ROS production.^47^ Therefore, one possible potential cofactors of NR2F2 is the hypoxia/ROS pathway. This new research direction will be our future focus, and it will identify the cofactors in CM.

This study has several limitations. First, WTA data were obtained from single blood vessels containing a mixture of ECs and SMCs. Data generated was not from a single cell resolution, which impedes further analysis of cell-cell interactions. Second, GeoMx is a hybridization-based assay. It is less sensitive but less biased than PCR-based RNA-seq platforms. Therefore, it may fail to identify some genes that are dysregulated in CM lesions. Third, the NR2F2-mediated pathways that lead to the EndMT acquisition of CM ECs have not yet been identified. Fourth, the structural, dynamic, and functional aberrances of AJs, their interactions with NR2F2, and their roles in EndMT in CM lesions have not been explored. These will be the scope of our future investigations.

## Acknowledgement

We appreciate Dr. Raju Pillai and Dr. Lixin Yang in the Department of Pathology, City of Hope National Medical Center, Los Angeles for the assistance of GeoMx WTA experiments. We are very thankful for the support and assistance from the University of South Carolina School of Medicine Instrumentation Resource Core Facility (RRID:SCR_024955). The Galaxy server used for some calculations is partly funded by the German Federal Ministry of Education and Research BMBF grant 031 A538A de.NBI-RBC and the Ministry of Science, Research and the Arts Baden-Württemberg (MWK) within the framework of LIBIS/de.NBI Freiburg.

## Database deposition

The scRNA-seq omics data has been deposited into the NCBI GEO (GSE295372). The GeoMx data has been deposited into the GEO (GSE297100). Data is available to the public.

## Resource availability Lead contact

Further information and requests for resources and reagents should be directed to and will be fulfilled by the lead contacts, Wenbin Tan and Yunguan Wang.

## Materials availability

Unique/stable reagents generated in this study are available from the lead contact Wenbin Tan with an MTA.

## Data and code availability

The GeoMx WTA data, CM phosphoproteomic data, and iEC ATAC-seq data are deposited into the NCBI GEO or as supplemental materials that are available to the public.

The code is available from the lead contact Yunguan Wang.

## Author contributions

W.T. designed and supervised the study. Y.W. designed and supervised the bioinformatic analysis of the study. V.N., together with I.C., S.H., M.A., Y.R., L.P., performed GeoMx, IF, scRNA-seq, Western blot, phosphoproteomic, iPSC culture experiments. I.M. performed scRNA-seq, RGN, and KO simulation analyses. M.L.H., E.H.C., and H.I.F. performed patient consents and biopsy collections. A.G.J. performed phosphoproteomic analysis. Y.C.C., Q.L., D.F., C.F.W., D.C., J.S.N., A.G.J., and Y.W. contributed to data analysis, data interpretation, and manuscript revision. V.N. and W.T. drafted the manuscript with input from all authors.

## Supplemental information

Supplemental method and discussion

Supplemental Figures 1-26

Supplemental tables 1-3

