## Supplementary material for "Single-vessel transcriptome map pathological landscapes and reveal NR2F2-mediated smooth muscle cell phenotype acquisition in capillary malformations": Suppl figures 1-26

**Supplemental Figures 1-26**

CM vs normal vessels_overall

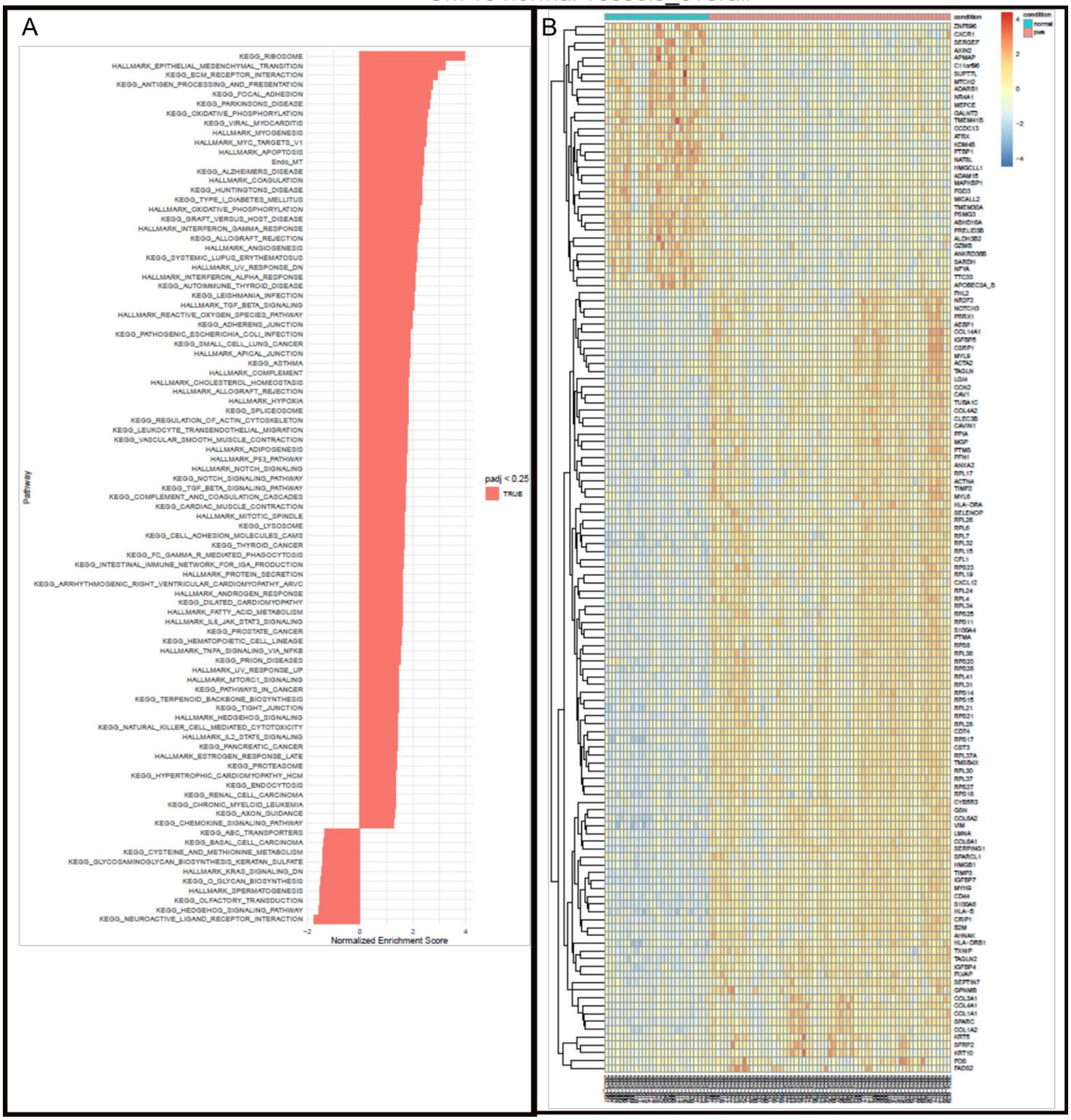


**Suppl figure 1:** (A) GSEA and (B) heatmap for CM vs CTL blood vessels by GeoMx WTA profiling.

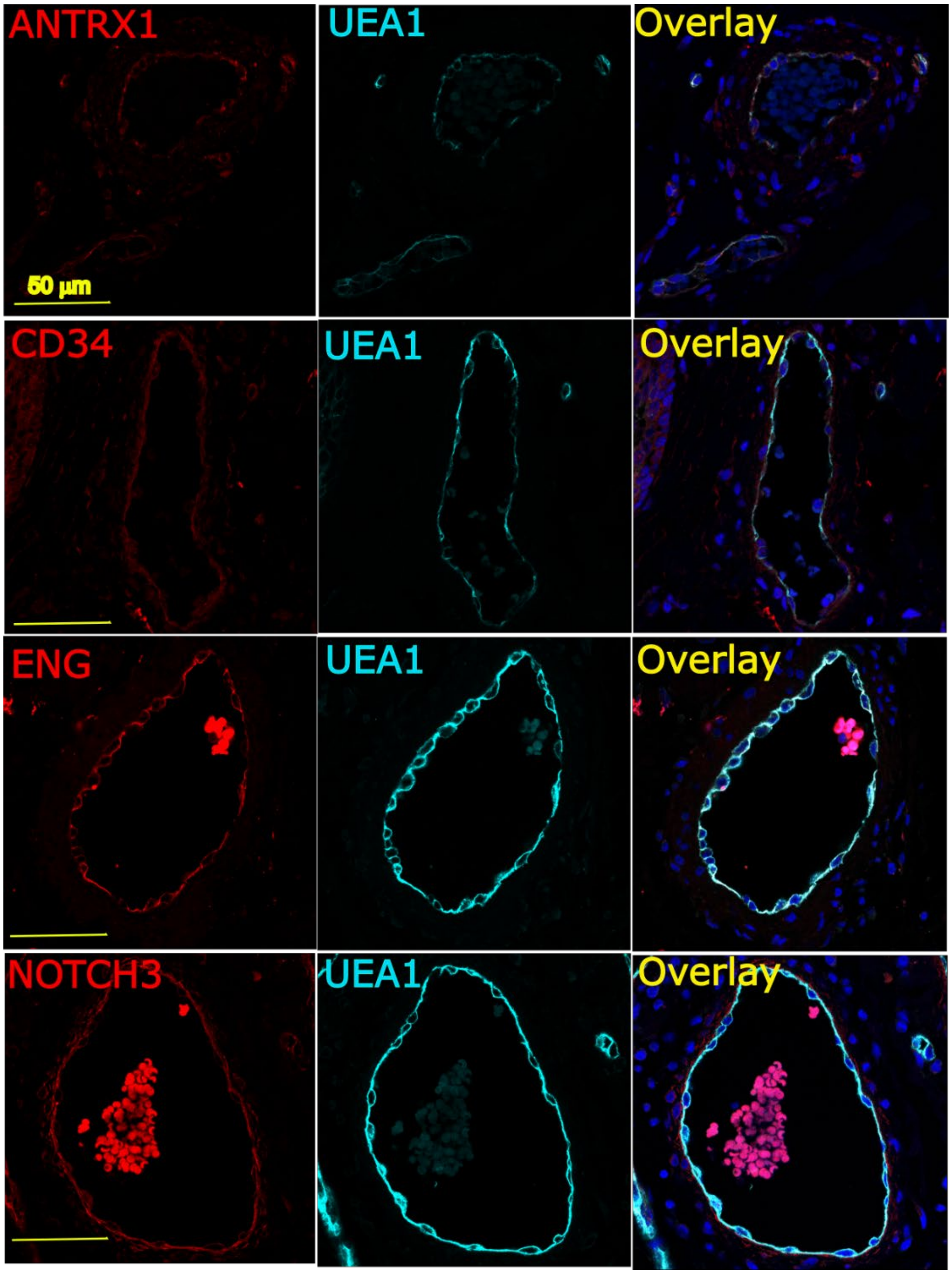


**Suppl figure 2:** IF validation of the expression patterns of DEGs related to EC or EPC biomarkers in CM vessels. Double staining of UEA1 (cyan) together with ANTRX1 (red), CD34 (red), ENG (red), or NOTCH3 (red). DAPI, blue.

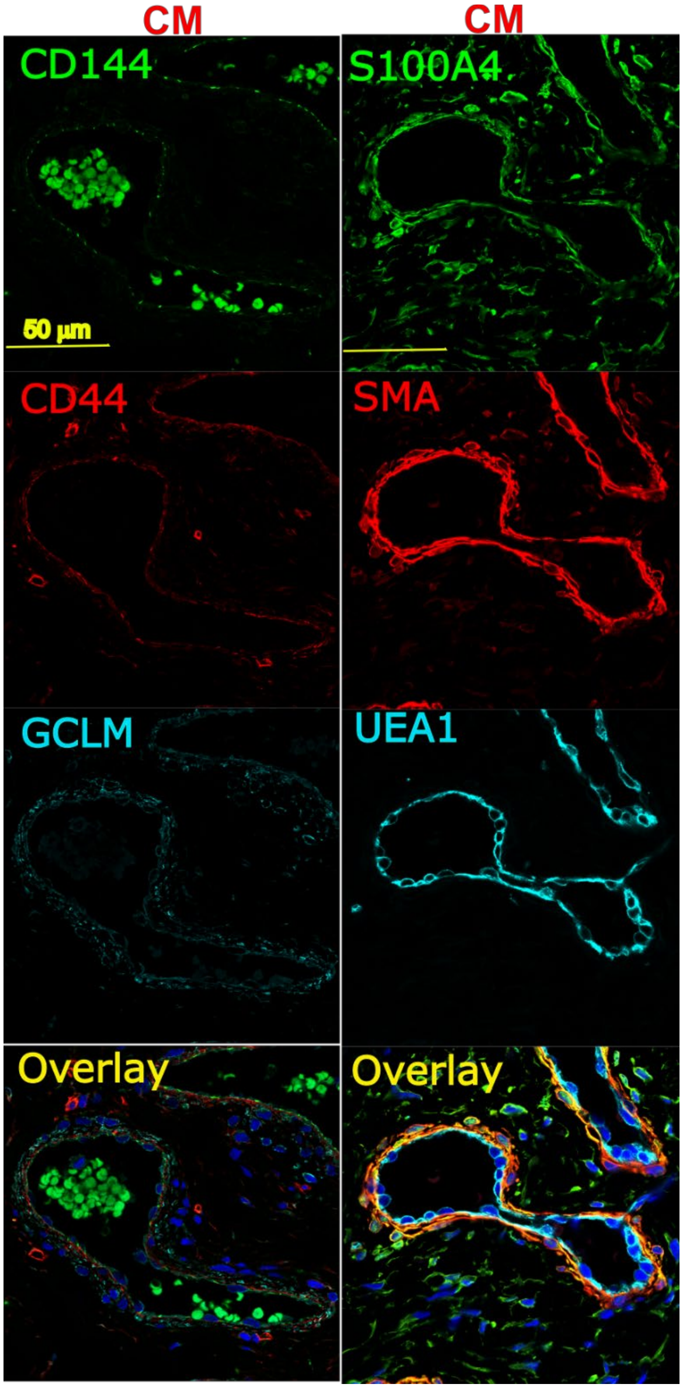


**Suppl figure 3:** IF validation of the expression patterns of EC-related DEGs including CD44 (red), GCLM (cyan), S100A4 (green), and SMA (red) in CM vessels. CD144 (green) or UEA1 (cyan) was used to show CM ECs. DAPI, blue.

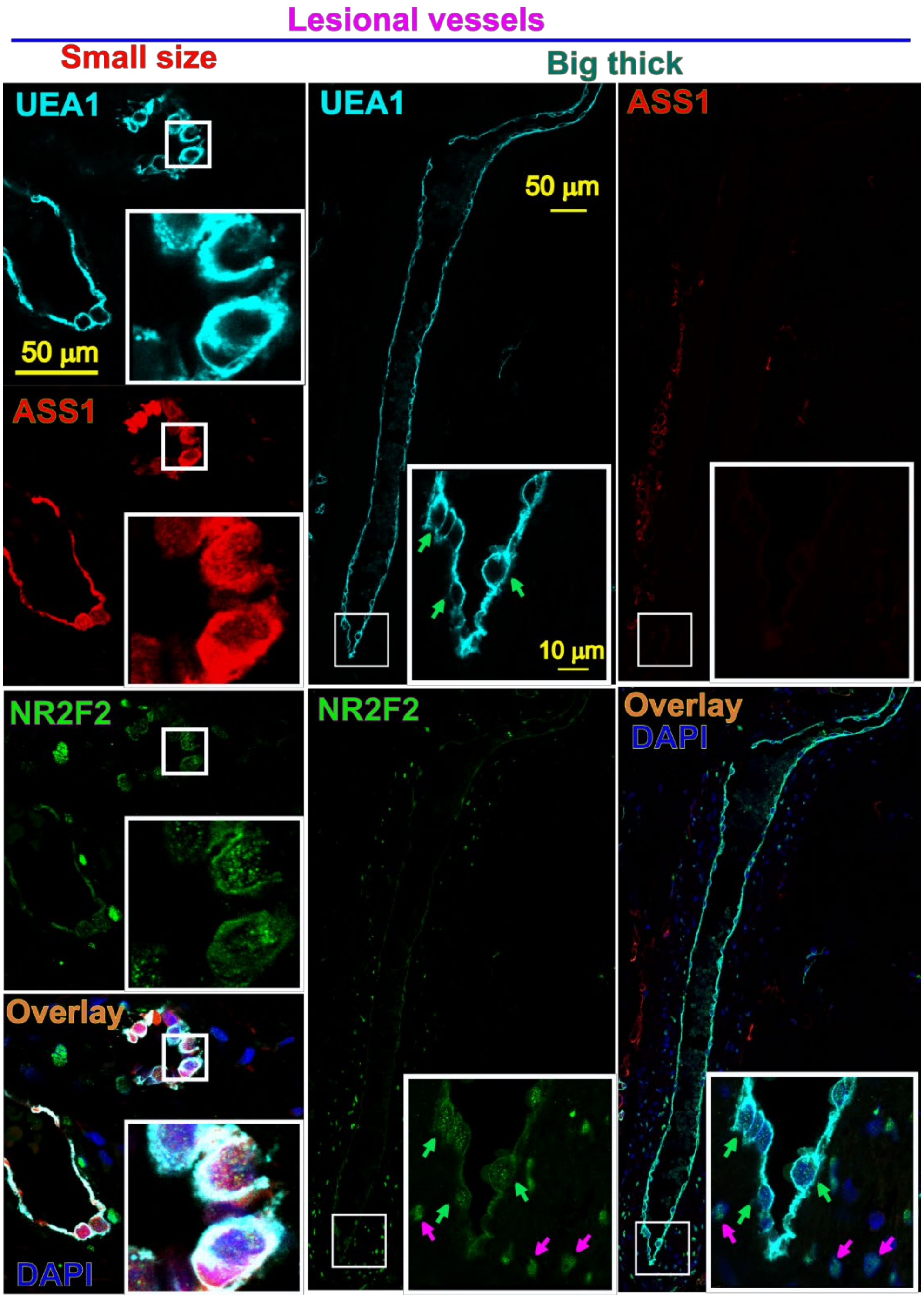


**Suppl figure 4:** The transiting patterns of NR2F2 and ASS1 in ECs of CM vessels with small or big sizes. Left-side panel: NR2F2 (green) and ASS1 (red) were found in the same ECs in small size CM blood vessels. Middle and right panels: in big thick lesional vessels, NR2F2 (green arrows) but not ASS1 was found in the nucleus of many ECs. Some scattered SMCs showed strong nucleus NR2F2 IF signaling. The right bottom insert was a high magnification of the box area from each image. UEA1 (cyan) staining was used to show ECs; DAPI: blue.

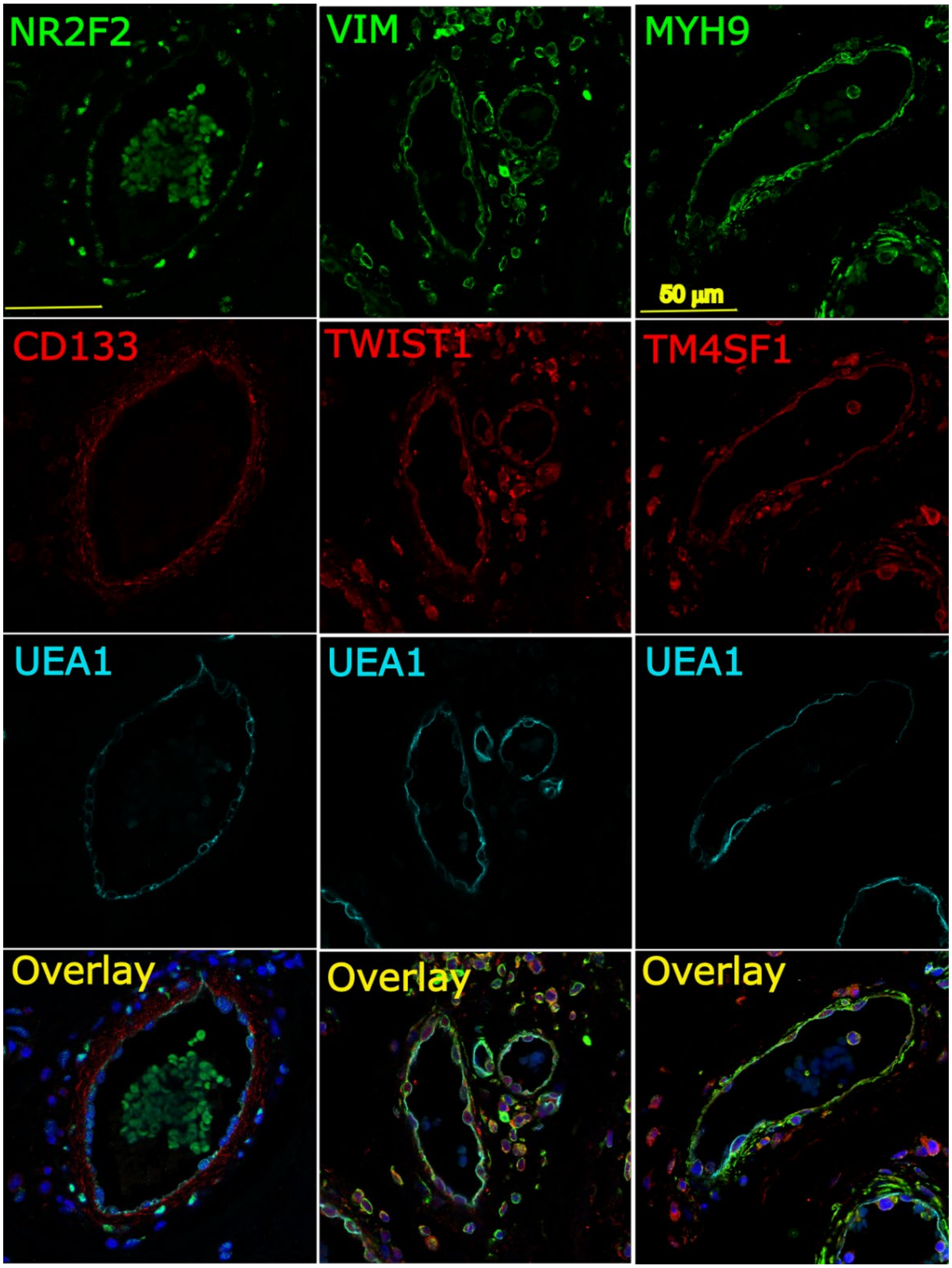


**Suppl figure 5:** IF validation of the expression patterns of CD133 and EndMT-related DEGs such as VIM, TWIST1, MYH9, and TM4SF1 in CM vessels. Triple staining of UEA1 (cyan) together with NR2F2 (green) plus CD133 (red), VIM (green) plus TWIST1 (red), or MYH9 (green) plus TM4SF1 (red). DAPI, blue.

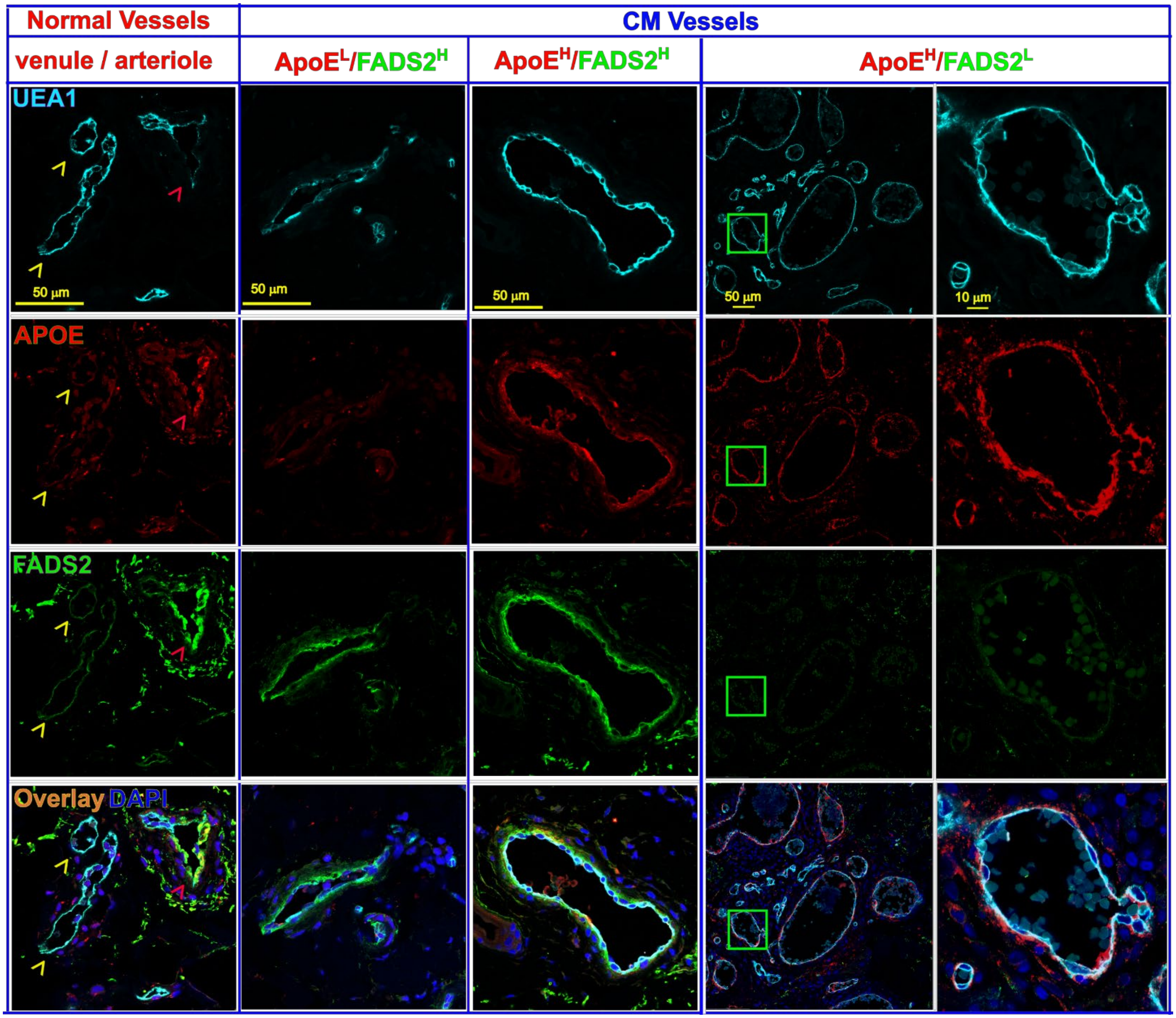


**Suppl figure 6:** Various expressing patterns of ApoE and FADS2 in CM vessels. In normal dermal vessels (left panel), FADS2 (green) but not ApoE (red) was found in ECs in capillaries and venules (yellow arrowheads); while both FADS2 and ApoE were highly present in arterioles (red arrowheads). The FADS2 was also found in the out layers of perivascular cells for arterioles. In CM vessels, three types of patterns were found, e.g., $ApoE^{L}/FADS2^{H}$, $ApoE^{H}/FADS2^{H}$, and $ApoE^{H}/FADS2^{L}$. The right-sided panel: a high magnification of green boxed areas from the image in the left. UEA1 (cyan) staining was used to show ECs; DAPI: blue.

Group 3_small_CM vs group 1_normal capillary_venule

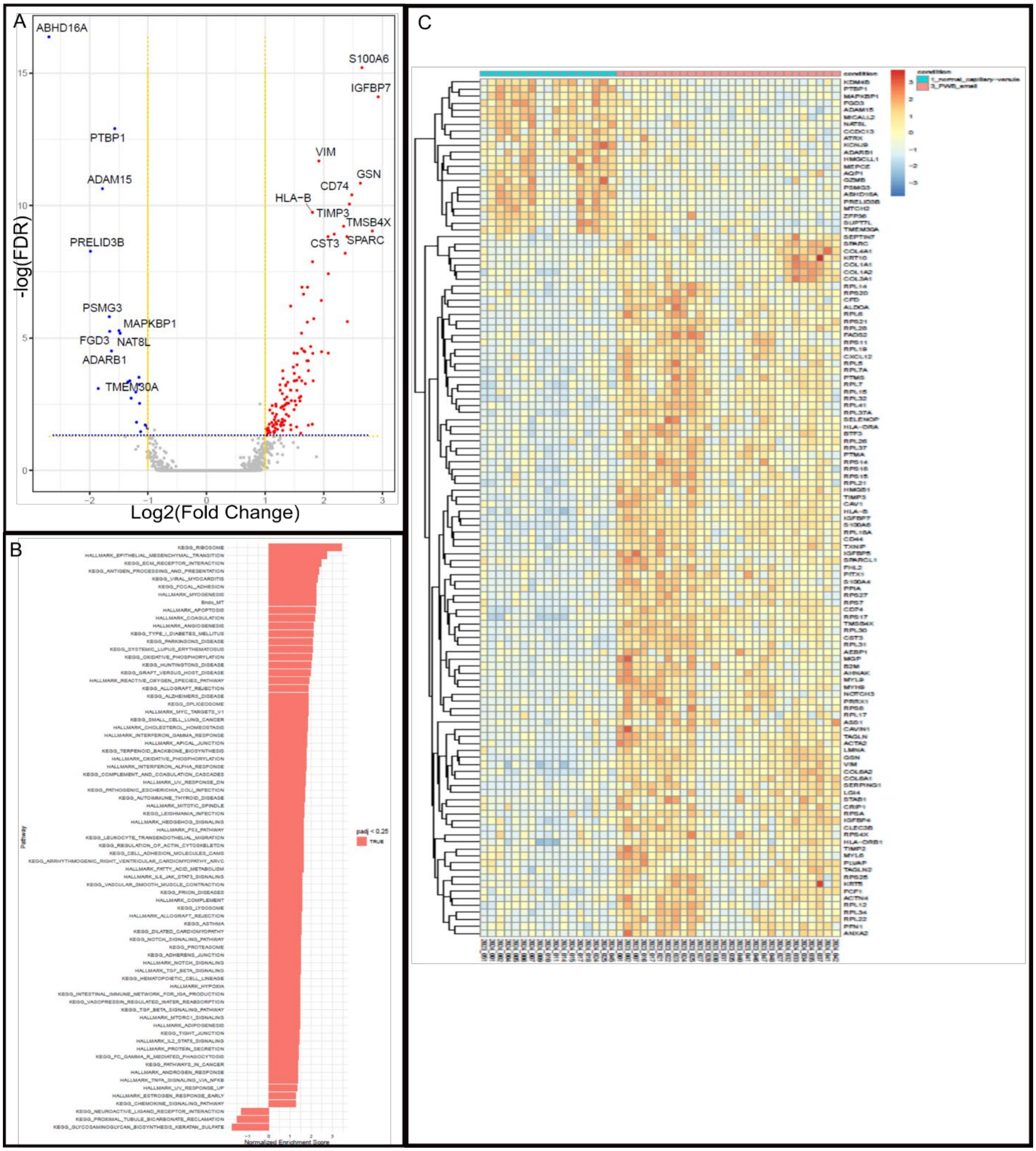


**Suppl figure 7**: (A) Volcano plot showing DEGs for small CM blood vessels (group 3) vs normal capillaries/venules (group 1); (B) GSEA and (C) heatmap for small CM blood vessels (group 3) vs normal capillaries/venules (group 1).

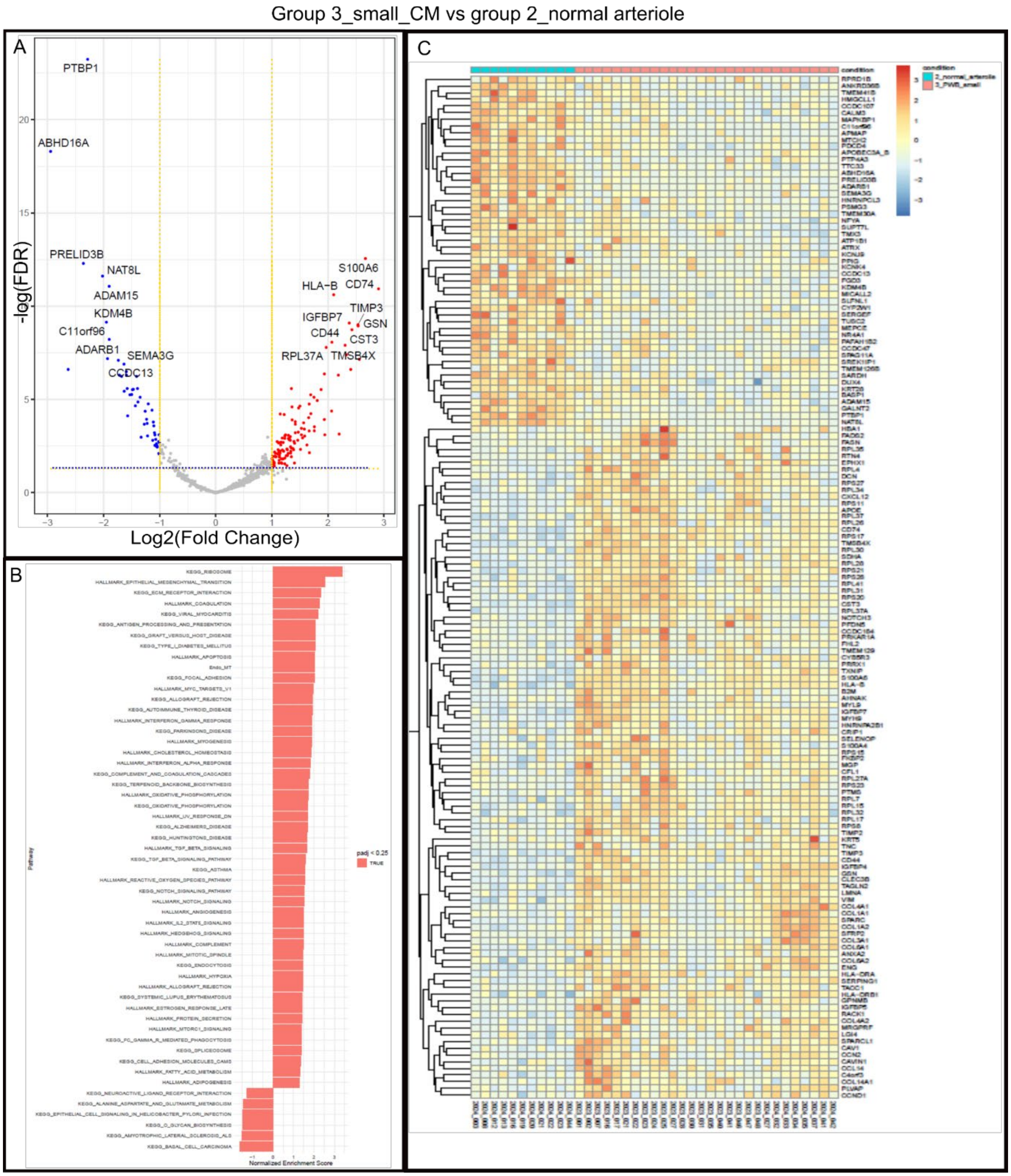


**Suppl figure 8**: (A) Volcano plot showing DEGs for small CM blood vessels (group 3) vs normal arterioles (group 2); (B) GSEA and (C) heatmap for small CM blood vessels (group 3) vs normal arterioles (group 2).

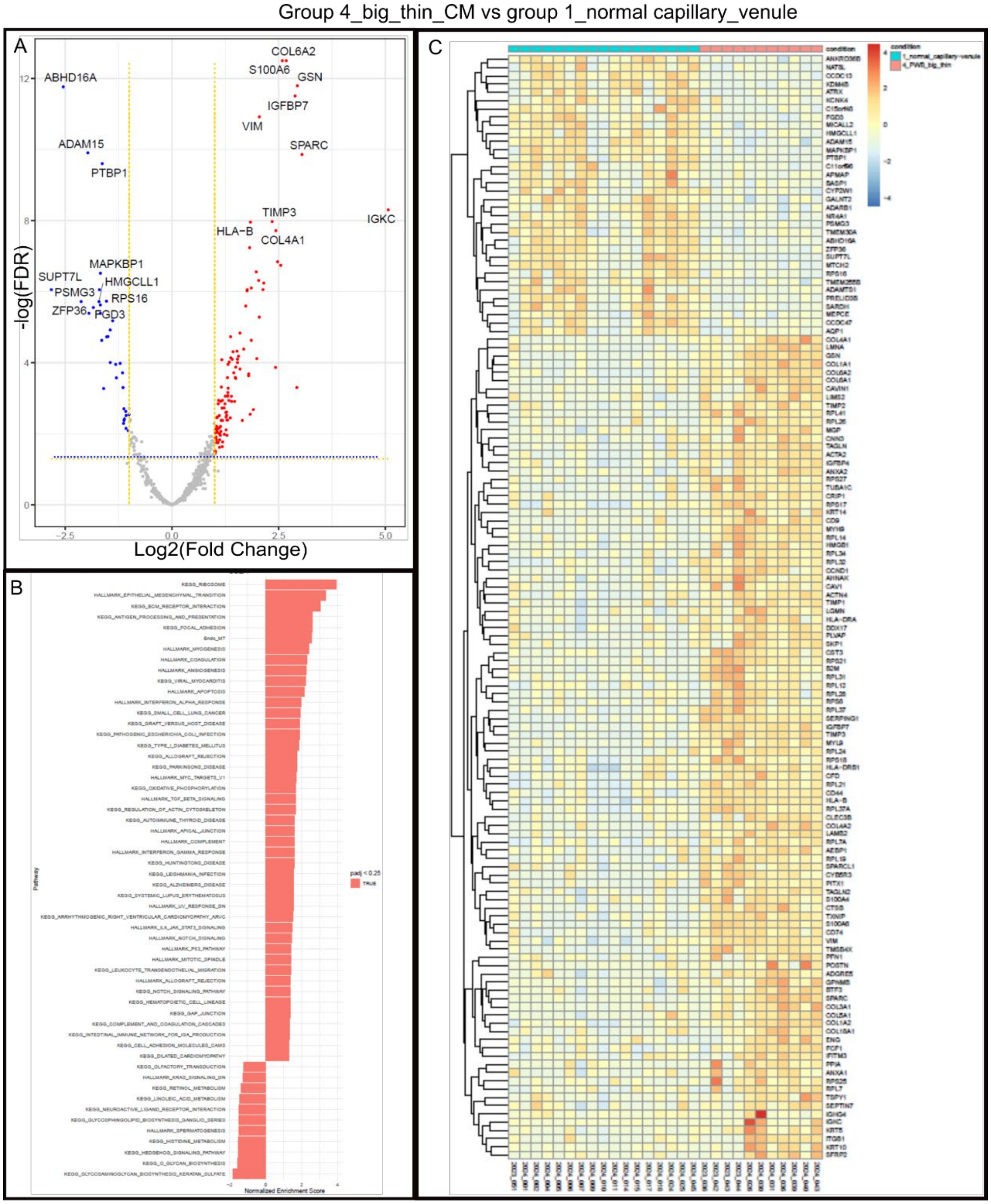


**Suppl figure 9:** (A) Volcano plot showing DEGs for big thin CM blood vessels (group 4) vs normal capillaries/venules (group 1); (B) GSEA and (C) heatmap for small CM blood vessels (group 3) vs normal capillaries/venules (group 1).

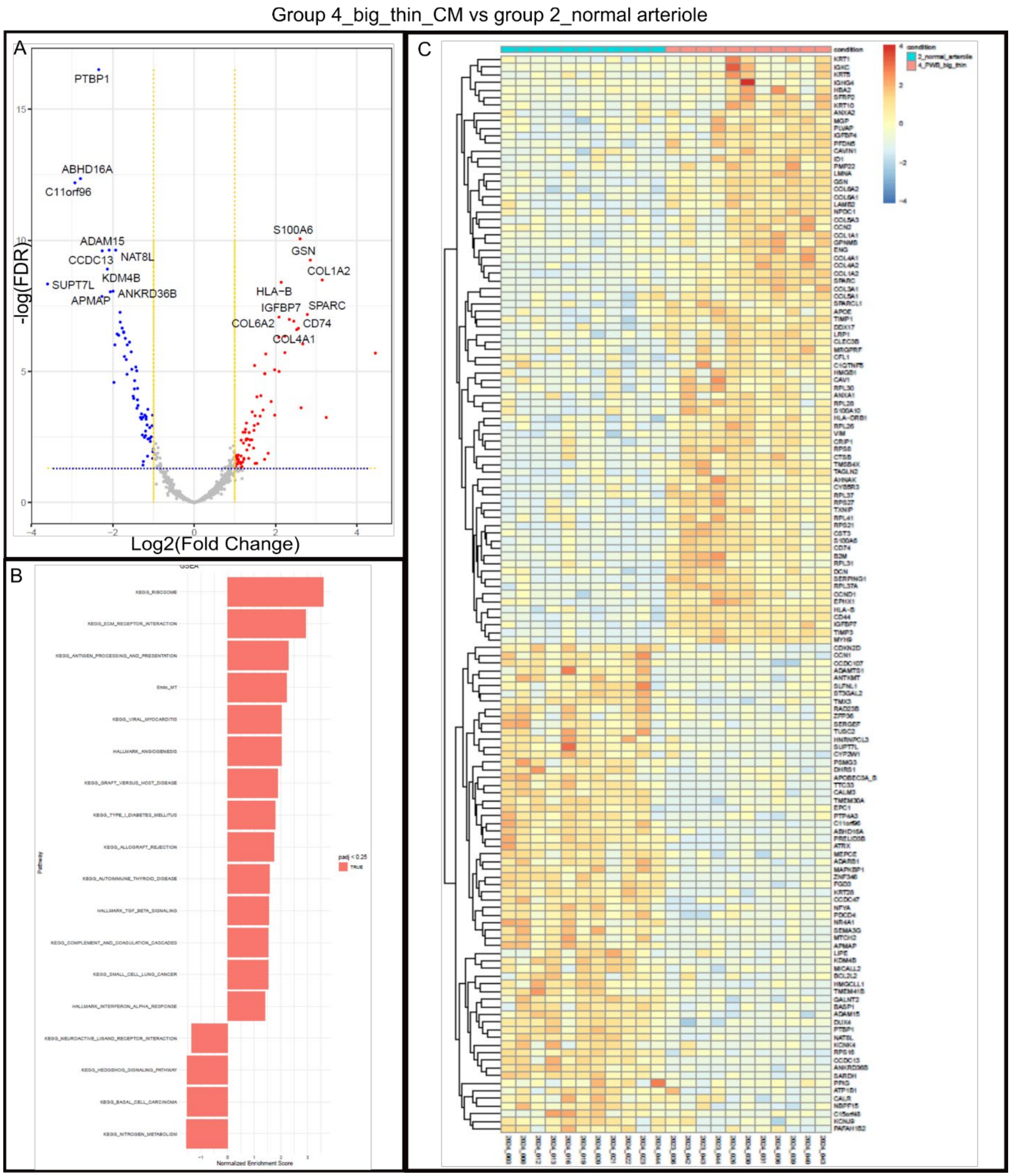


**Suppl figure 10:** (A) Volcano plot showing DEGs for big thin CM blood vessels (group 4) vs normal arterioles (group 2); (B) GSEA and (C) heatmap for small CM blood vessels (group 3) vs normal arterioles (group 2).

Group 5_big_thick_CM vs group 1_normal capillary_venule

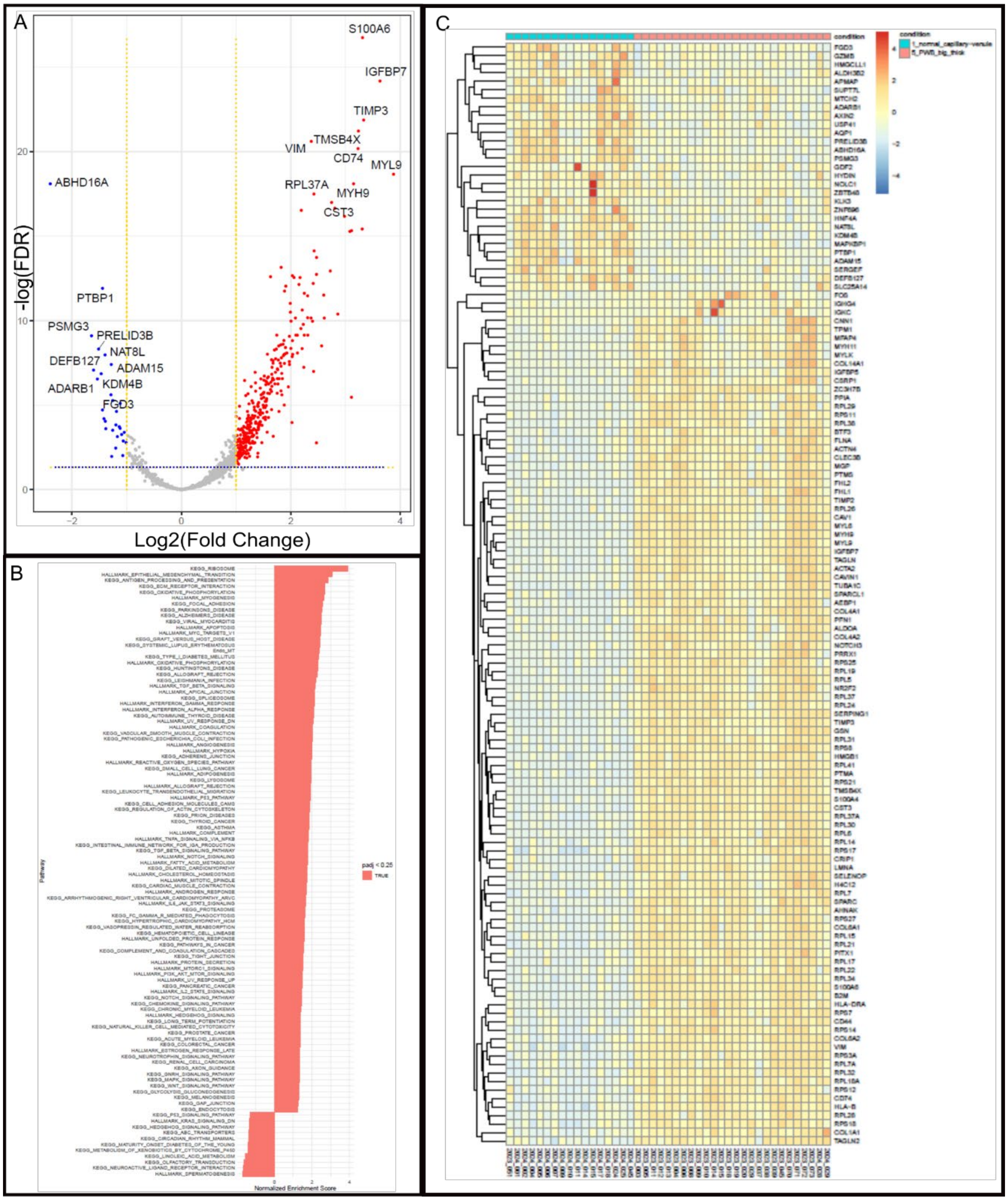


**Suppl figure 11:** (A) Volcano plot showing DEGs, (B) GSEA, and (C) heatmap for big thick CM blood vessels (group 5) vs normal capillaries/venules (group 1).

Group 5_big_thick_CM vs group 2_normal arteriole

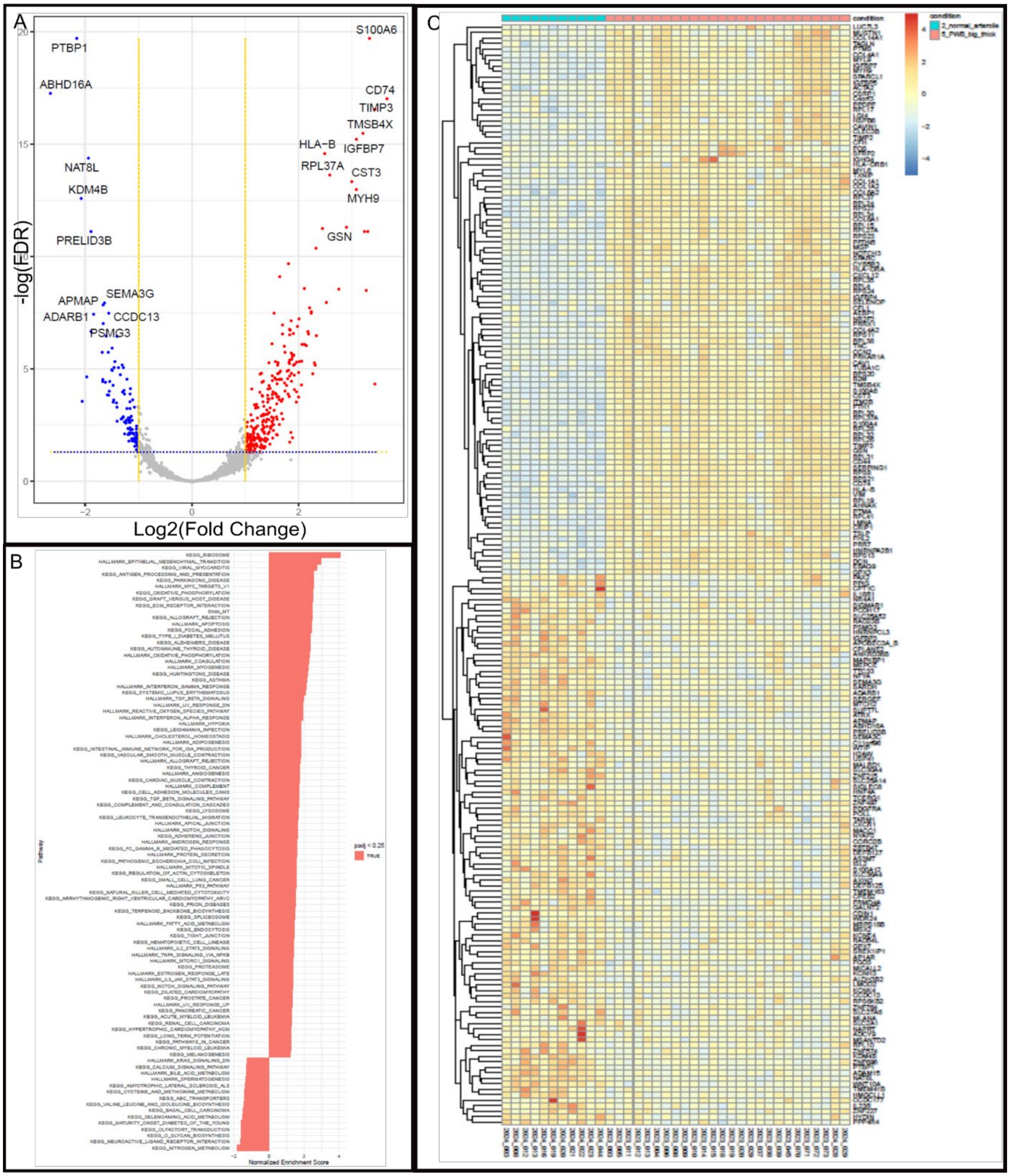


**Suppl figure 12:** (A) Volcano plot showing DEGs, (B) GSEA, and (C) heatmap for big thick CM blood vessels (group 5) vs normal arterioles (group 2).

Group 5_big_thick_CM vs group 3_small CM

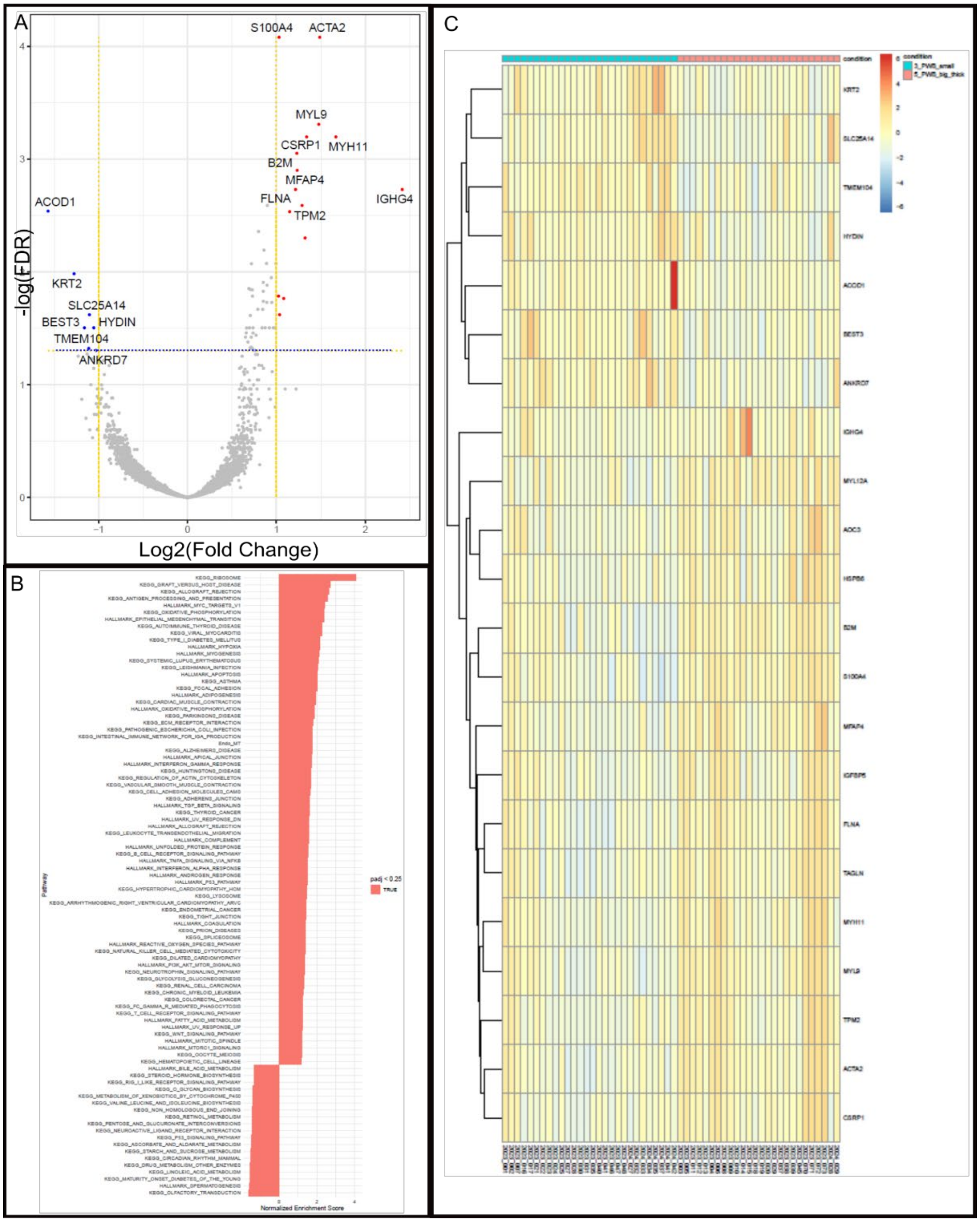


**Suppl figure 13:** (A) Volcano plot showing DEGs, (B) GSEA, and (C) heatmap for big thick (group 5) vs small CM blood vessels (group 3).

Group 5_big_thick_CM vs group 4_big_thin_CM

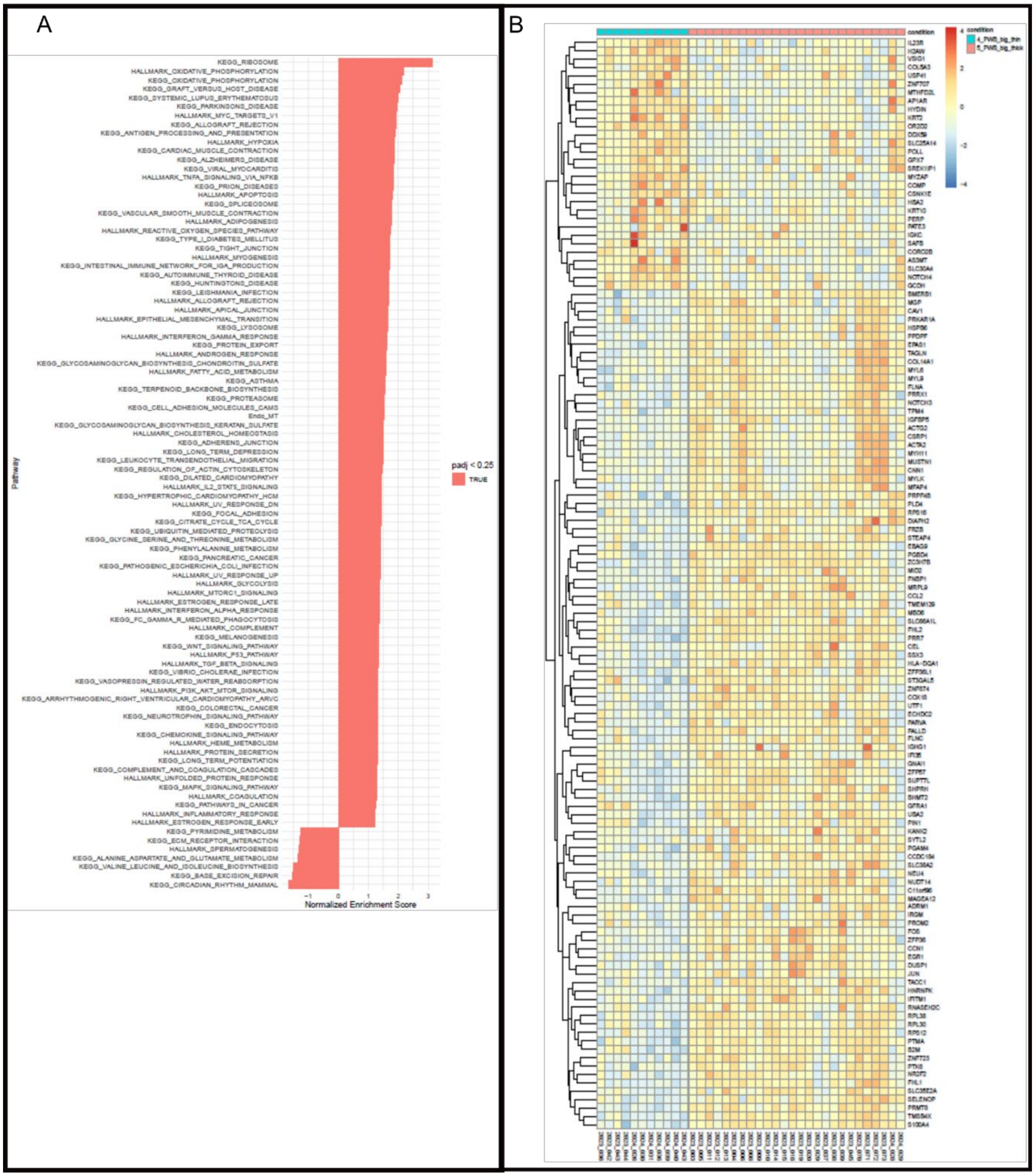

**Suppl figure 14:** (A) GSEA and (B) heatmap for big thick (group 5) vs big thin CM blood vessels (group 4) by GeoMx WTA profiling.

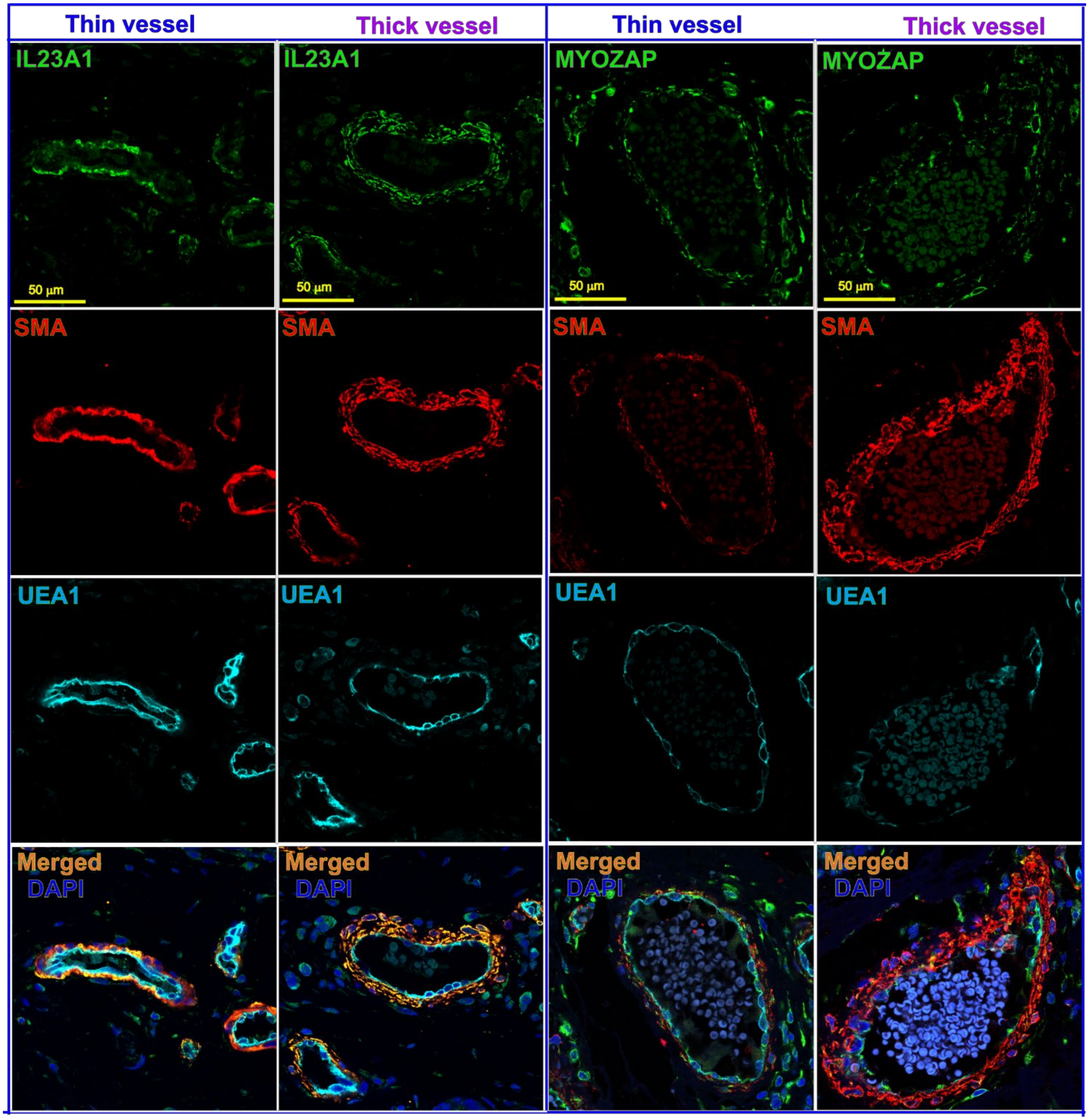


**Suppl figure 15:** IF validation for expression patterns of IL23A1 and MYOZAP in thin and thick CM vessels. Triple staining of IL23A1 (green), SMA (red), and UEA1 (cyan) showed that IL23A1 was mainly observed in SMCs but not ECs. The immunostaining signal intensity was similar in both types of big thin and thick CM vessels. Triple staining of MYOZAP (green), SMA (red), and UEA1 (cyan) showed that MYOZAP was observed in both ECs and SMCs. It exhibited punctate, fragmented and discorded immunostaining patterns with less intensity in big thick CM vessels than big thin ones.

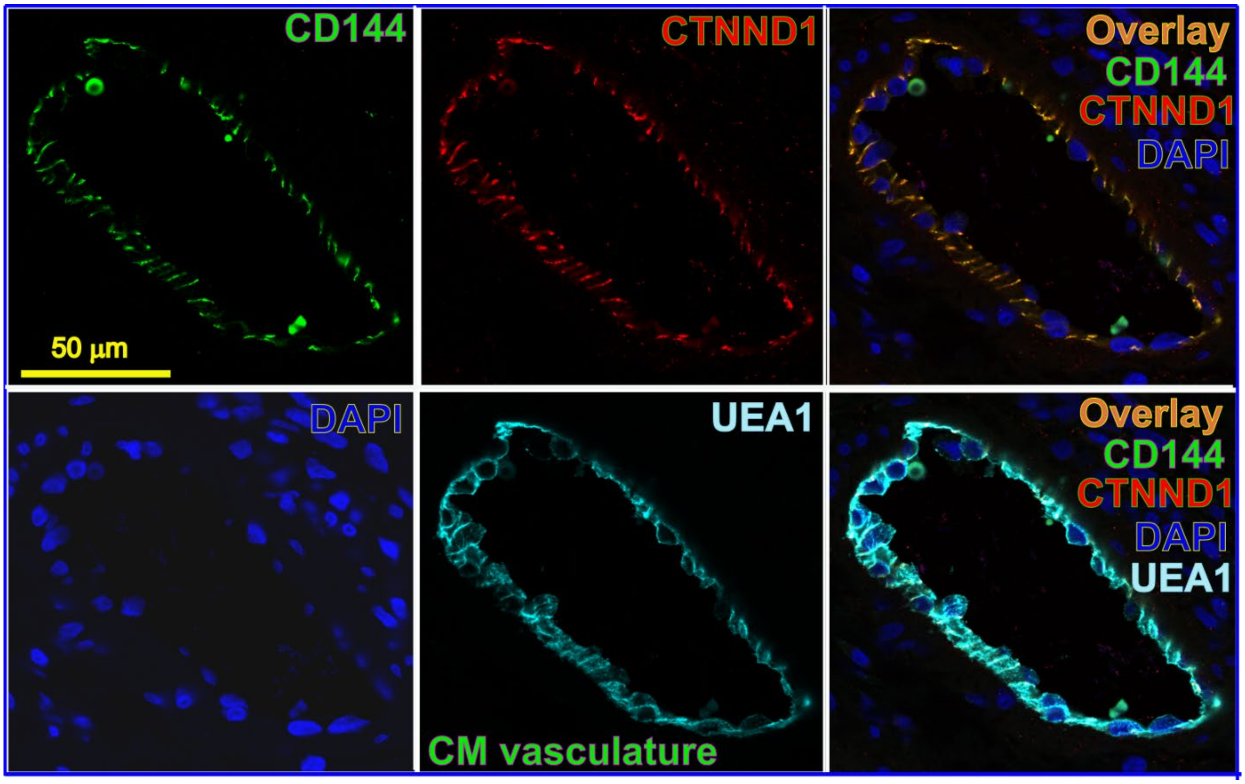


**Suppl figure 16:** IF validation for colocalization of CTNND1 and CDH5 among ECs in CM vessels. Triple staining of CDH5 (a.k.a., CD144, green), CTNND1 (red), and UEA1 (cyan) showing the colocalization of CDH5 and CTNND1 at inter-endothelial junctional sites.

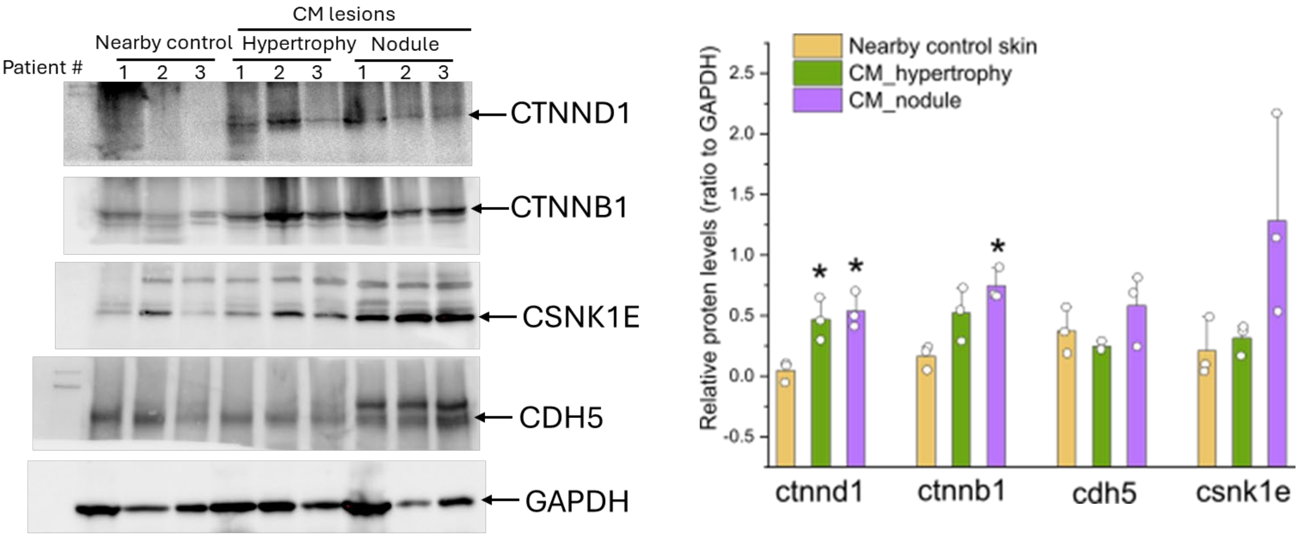


**Suppl figure 17:** The total protein levels of CTNND1, CTNNB1, CSNK1E, and CDH5 in CM lesions. Left panel: proteins were extracted from hypertrophic, nodular CM lesions and nearby normal control skins to CM lesions from the same patients (n=3), followed by Western blot analyses. Right panel: relative protein levels of CTNND1, CTNNB1, CSNK1E, and CDH5 to GAPDH. * $p<0.05$ comparing to nearby controls, paired *t*-test.

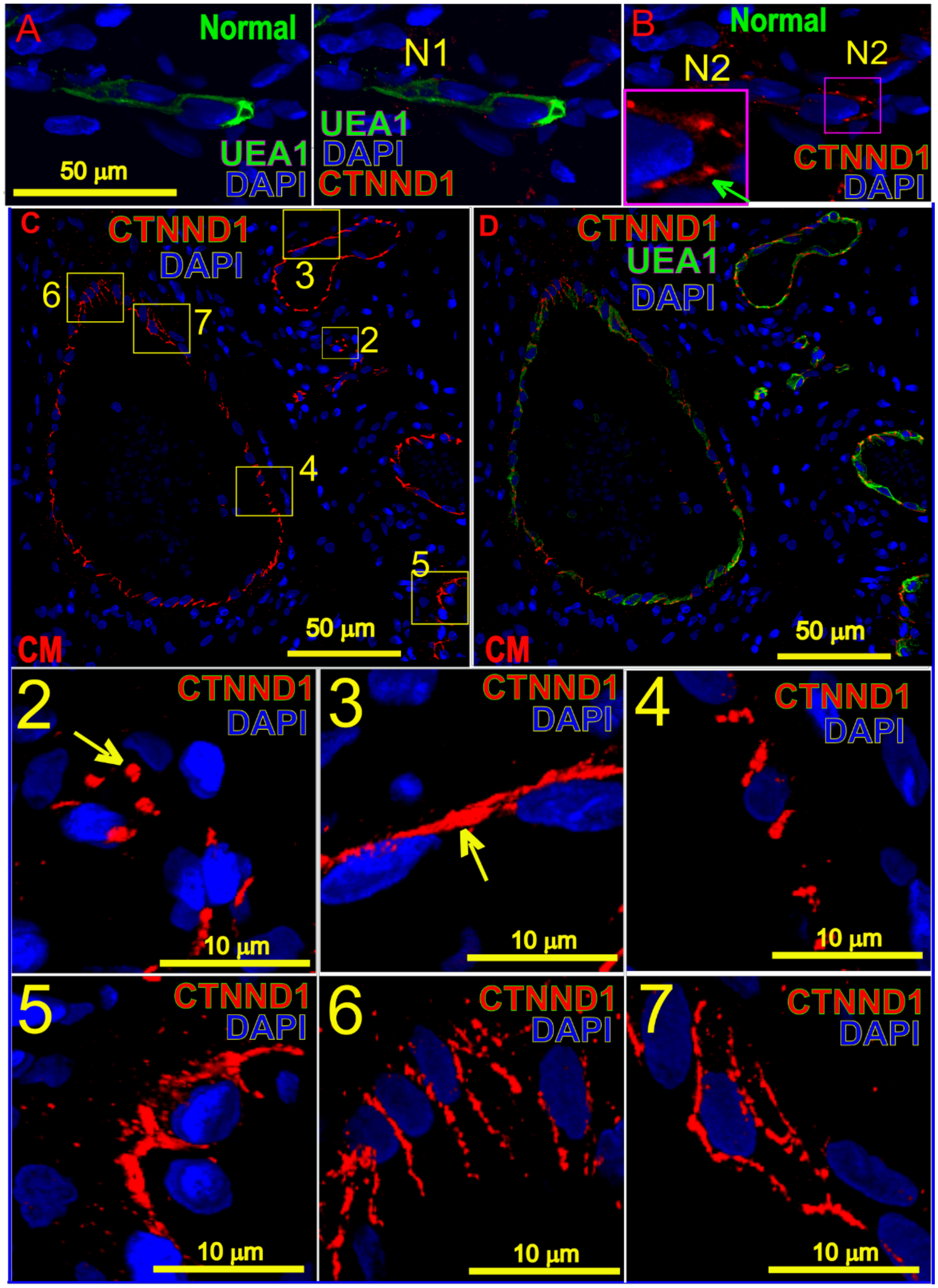


**Suppl figure 18:** Heterogenous endothelial CTNND1 patterns during CM EC phenotypic remodeling. (A) and (B) Normal human dermal capillary ECs showing either barely detectable CTNND1 signals (N1) or (B) CTNND1$^{pn}$ (N2). (C) and (D) CM ECs represent various patterns of CTNND1 among lesional blood vessels. Numbered boxed areas in (C) and (D) are magnified and shown in the individual panels with the same number, showing the heterogenous types of CTNND1 patterns; UEA1 (green) staining was used to show the morphologies of vasculature. Z-stacks confocal images were processed for CTNND1 subcellular patterns.

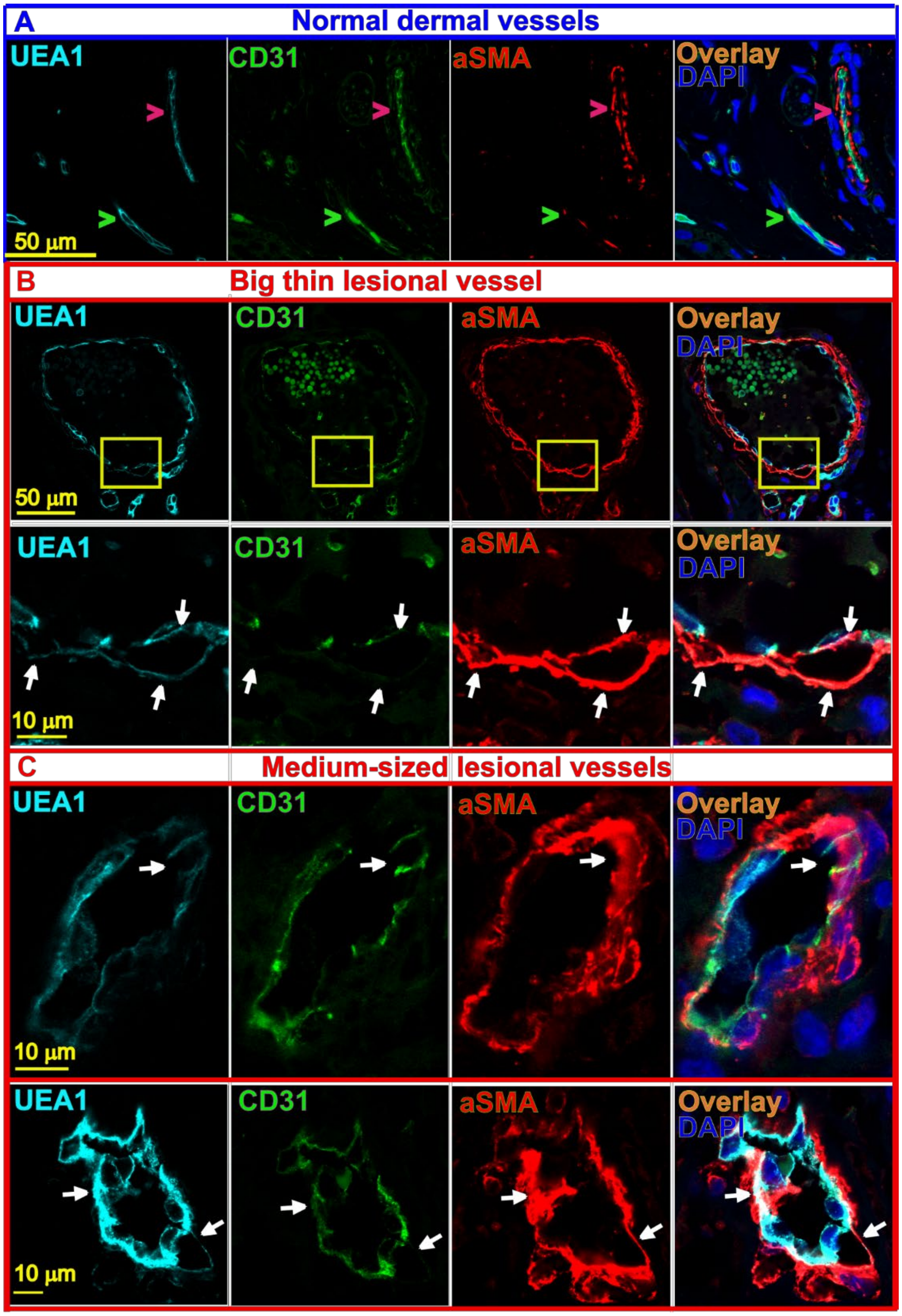


**Suppl figure 19:** CM ECs undergo EndMT remodeling. (A) A normal human dermal vessel with IF staining using antibodies against aSMA (red), UEA1 (cyan), and CD31 (green); (B) A big thin CM lesional vessel stained with the same antibodies. Lower panel: a high magnification of the boxed area (yellow box) from the upper panel showing an EndMT zone with two remodeling ECs (UEA1, blue) losing part of CD31 (green) signal but acquisition of SMA signal (white arrowheads). (C) Two medium-sized CM blood vessels showing the remodeling ECs (white arrowheads) with acquisition of SMA phenotype. UEA1 (cyan), CTNND1 (green), SMA (red), and DAPI (blue).

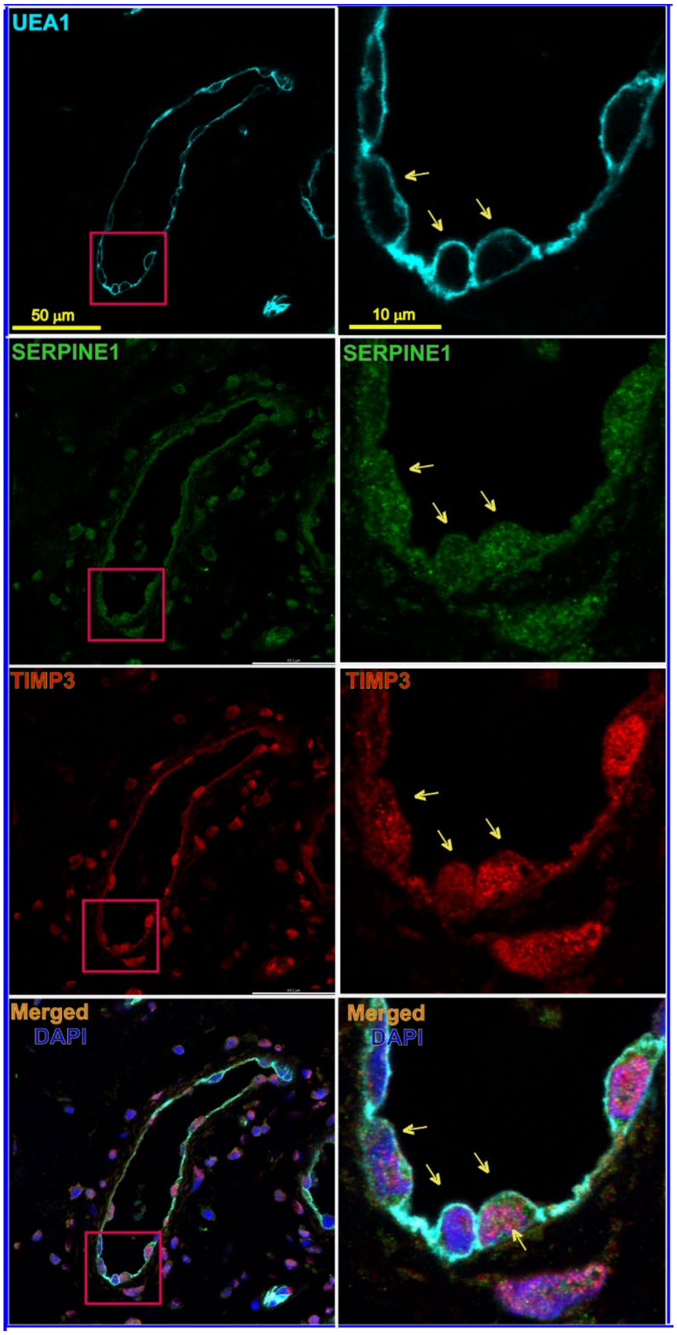


**Suppl figure 20:** IF validation of the expression patterns of DEGs including SERPINE 1 (green) and TIMP3 (red) in CM vessels. UEA1 (cyan) was used to show CM ECs. DAPI, blue. Right panel: a high magnification of red box area from the left panel. ECs in CM vessels showed co-expression of SERPINE 1 and TIMP3 (yellow arrowheads).

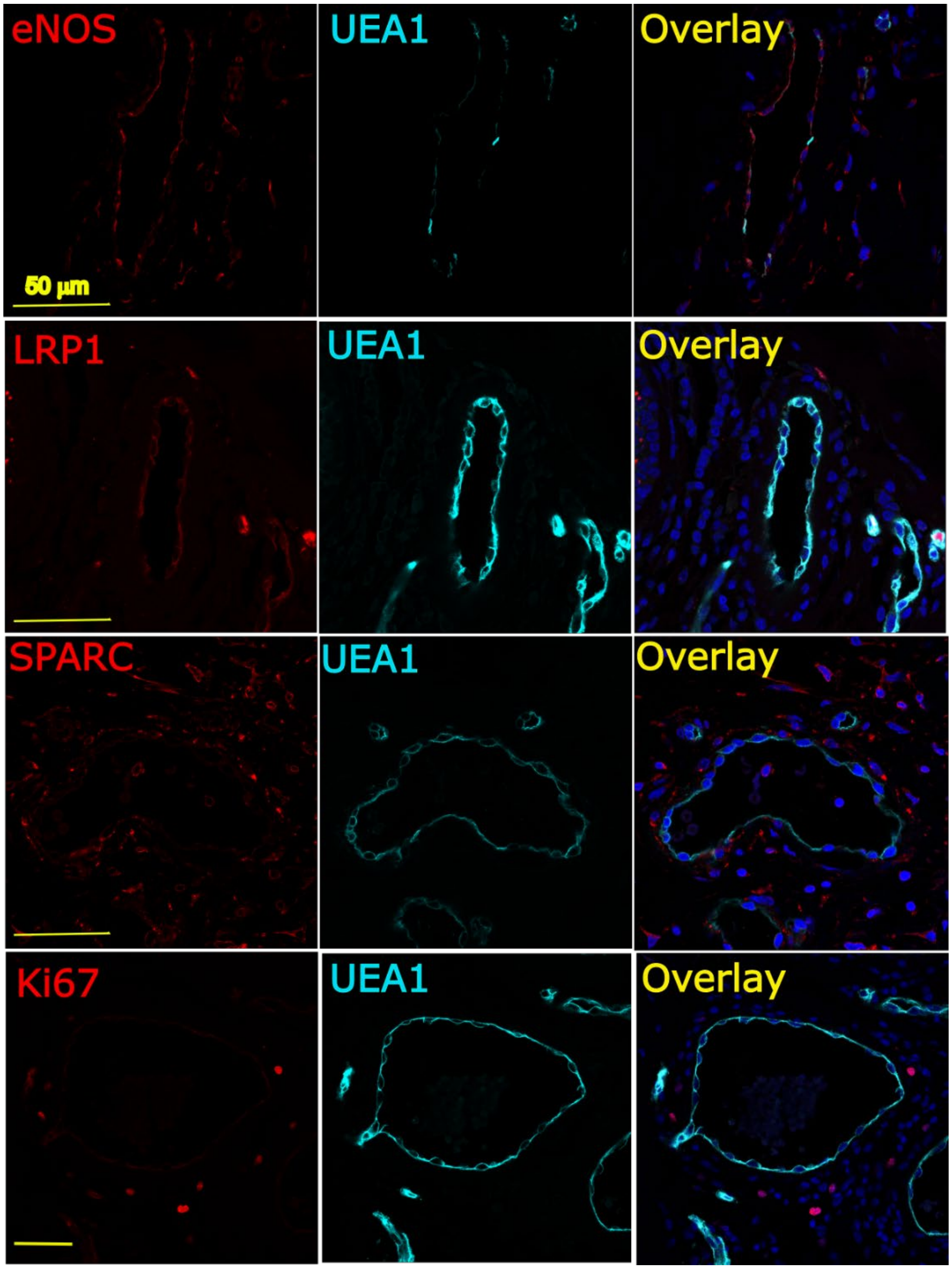


**Suppl figure 21:** IF validation of the expression patterns of metabolic remodeling related DEGs including eNOS (red), LRP1 (red), and SPARC (red) in CM vessels. UEA1 (cyan) was used to show CM ECs. DAPI, blue. Very few scattered Ki67 positive nuclei could be found among perivascular cells. No ECs with Ki67 positive nuclei were found in CM vessels.

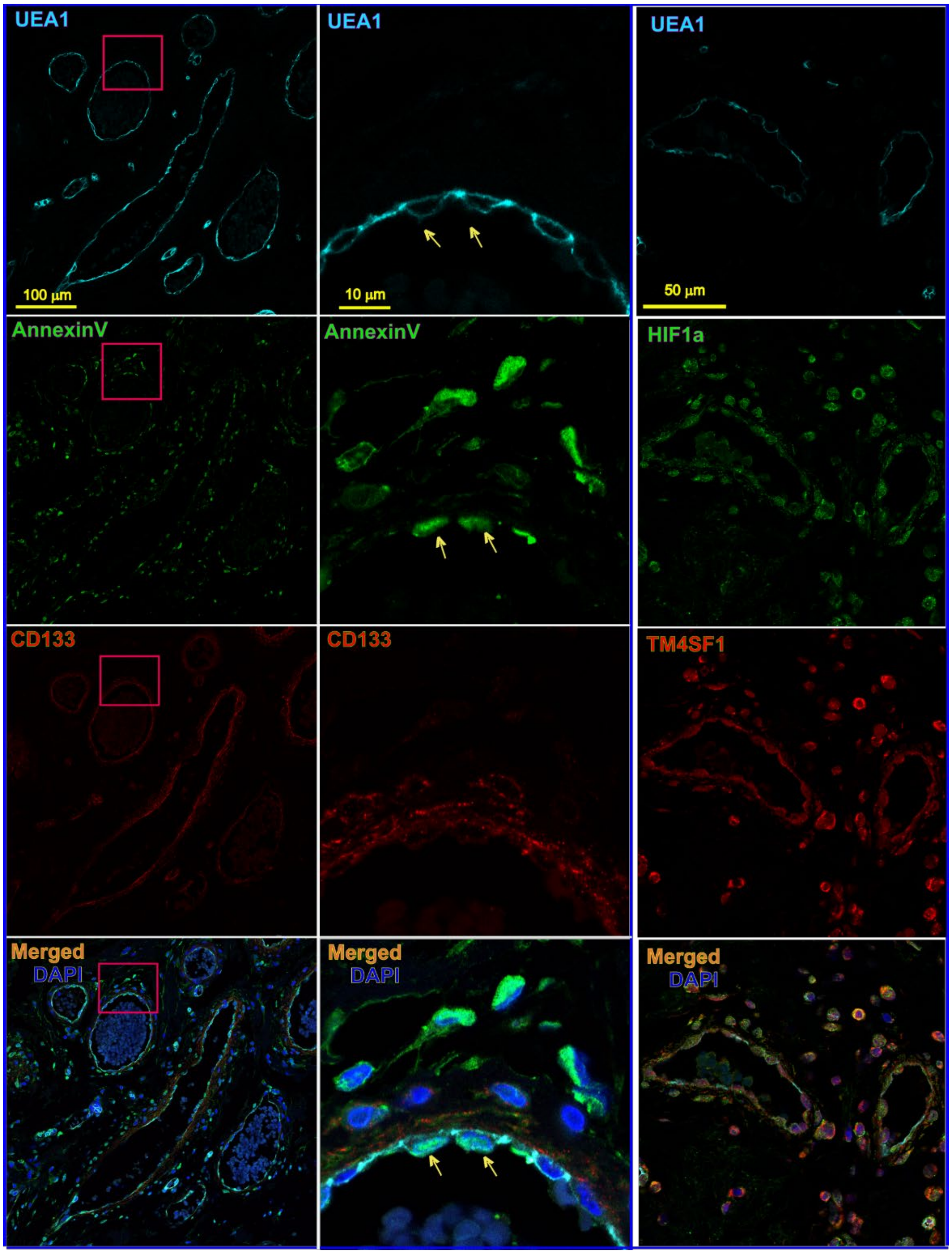


**Suppl figure 22:** IF validation of the expression patterns of DEGs including Annexin V and HIF1a in CM vessels. Left panel: triple IF staining of UEA1 (cyan), Annexin V (green), and CD133 (red). Middle panel: a high magnification of red box area from the left panel. ECs in CM vessels showed co-expression of Annexin V and membrane CD133 (yellow arrowheads). Some scattered Annexin V perivascular cells were observed. Right panel: triple IF staining of UEA1 (cyan), HIF1a (green), and TM4SF1 (red). ECs and many perivascular cells showed positive for both HIF1a and TM4SF1. DAPI, blue.

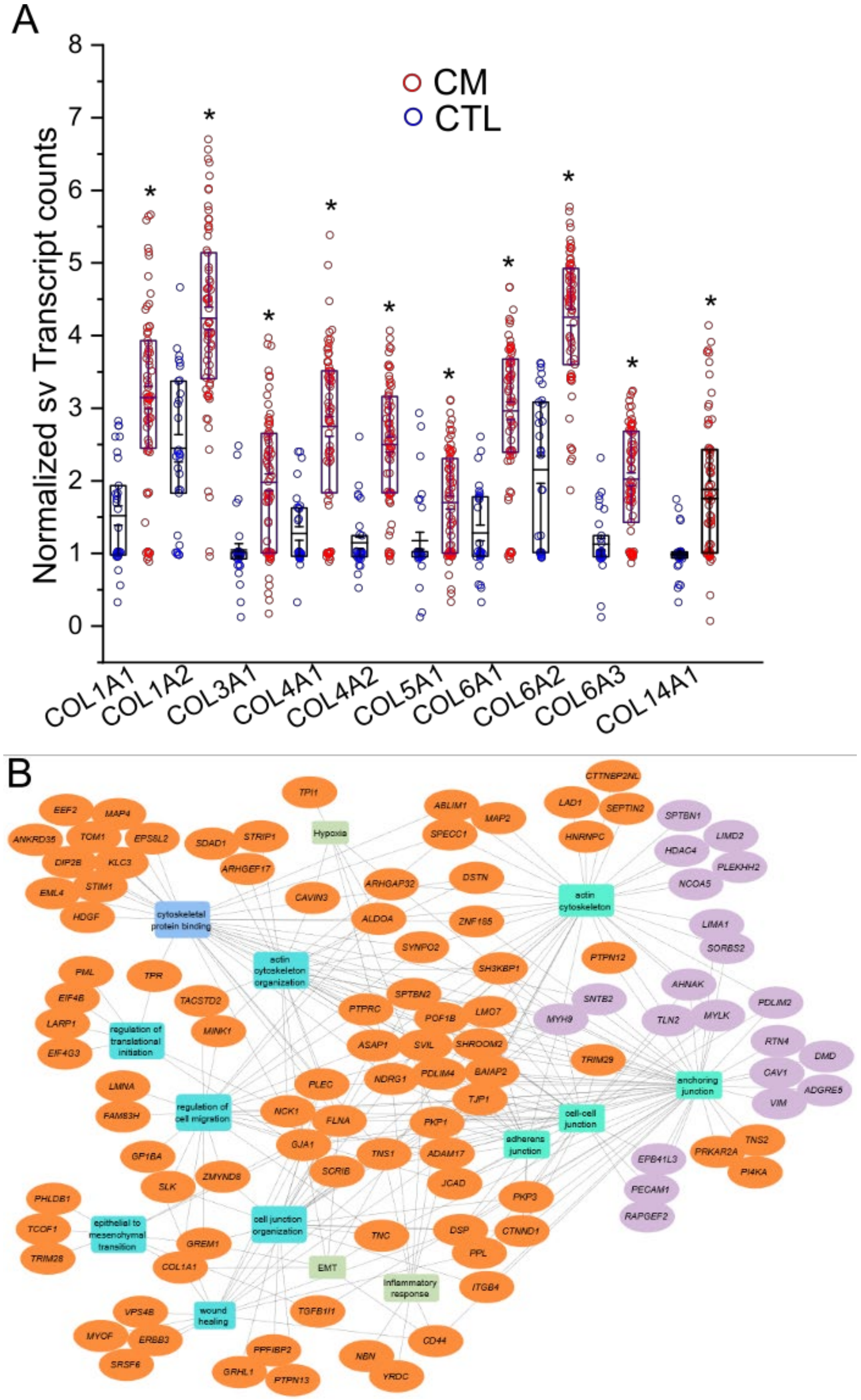


**Suppl figure 23:** (A) Representative DEGs of collagen profiles in CM vs normal dermal vessels. * FDR < 0.05. Each dot represents a single vessel. (B) DEP functional map showing DEPs and their enriched pathways involving anchoring and cell-cell junctions, AJs, Rho GTPases, hypoxia signaling, epithelial mesenchymal transition, regulation of cell migration, inflammatory response, and cytoskeleton organization. # $p<0.05$.

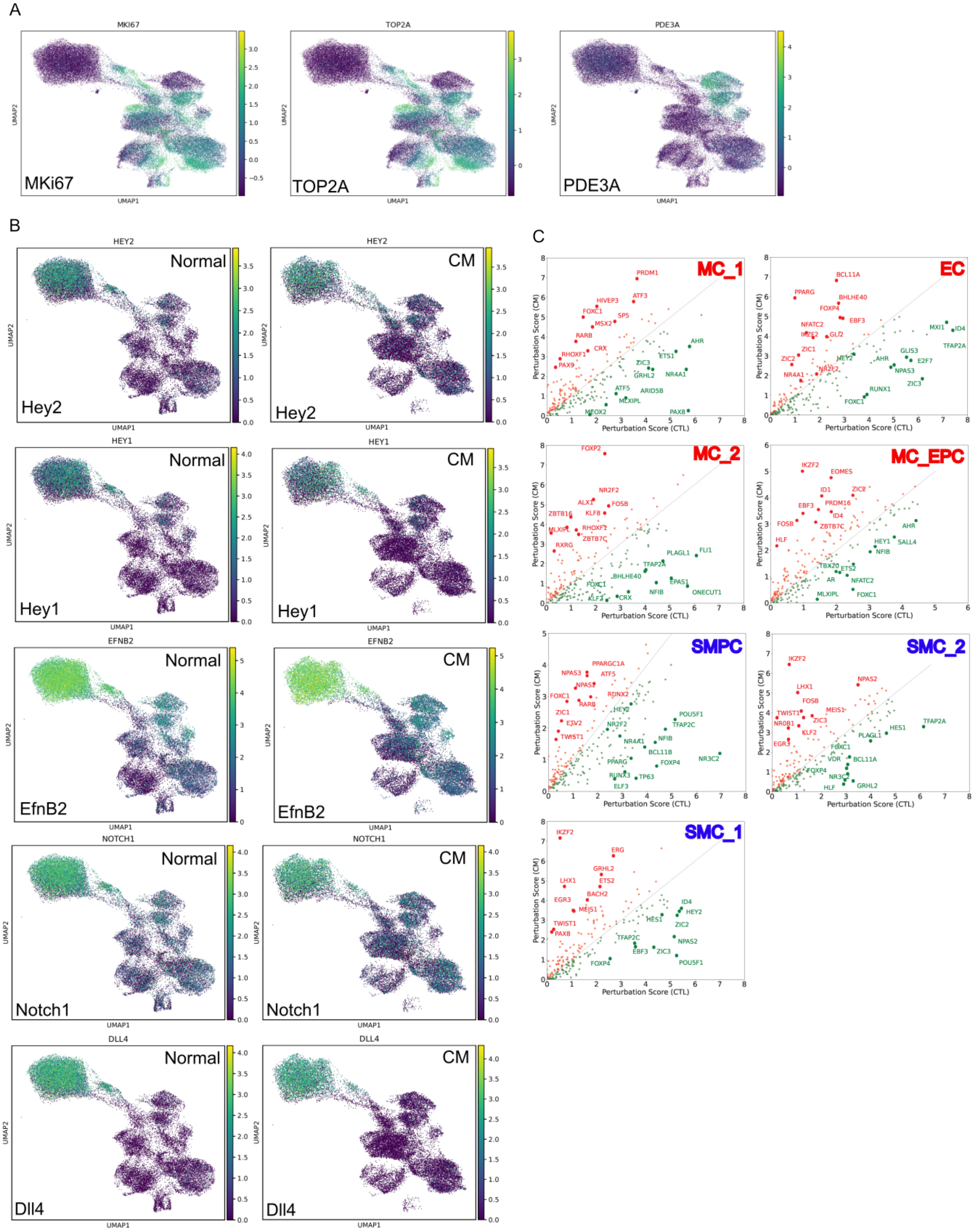


**Suppl figure 24** (A) Expressions of MKi67, TOP2A, and PDE3A among clusters. (B) Attenuation of expression of arterial biomarkers of Hey2, Hey1, EfnB2, Notch1, and Dll4 in EC cluster in CM as compared with CTL. (C), Systemic KO simulation result of top TFs with greater (red color labelled TFs) or less impacts (green color labelled TFs) in cell identity shifts in major clusters in CM as compared to CTL.

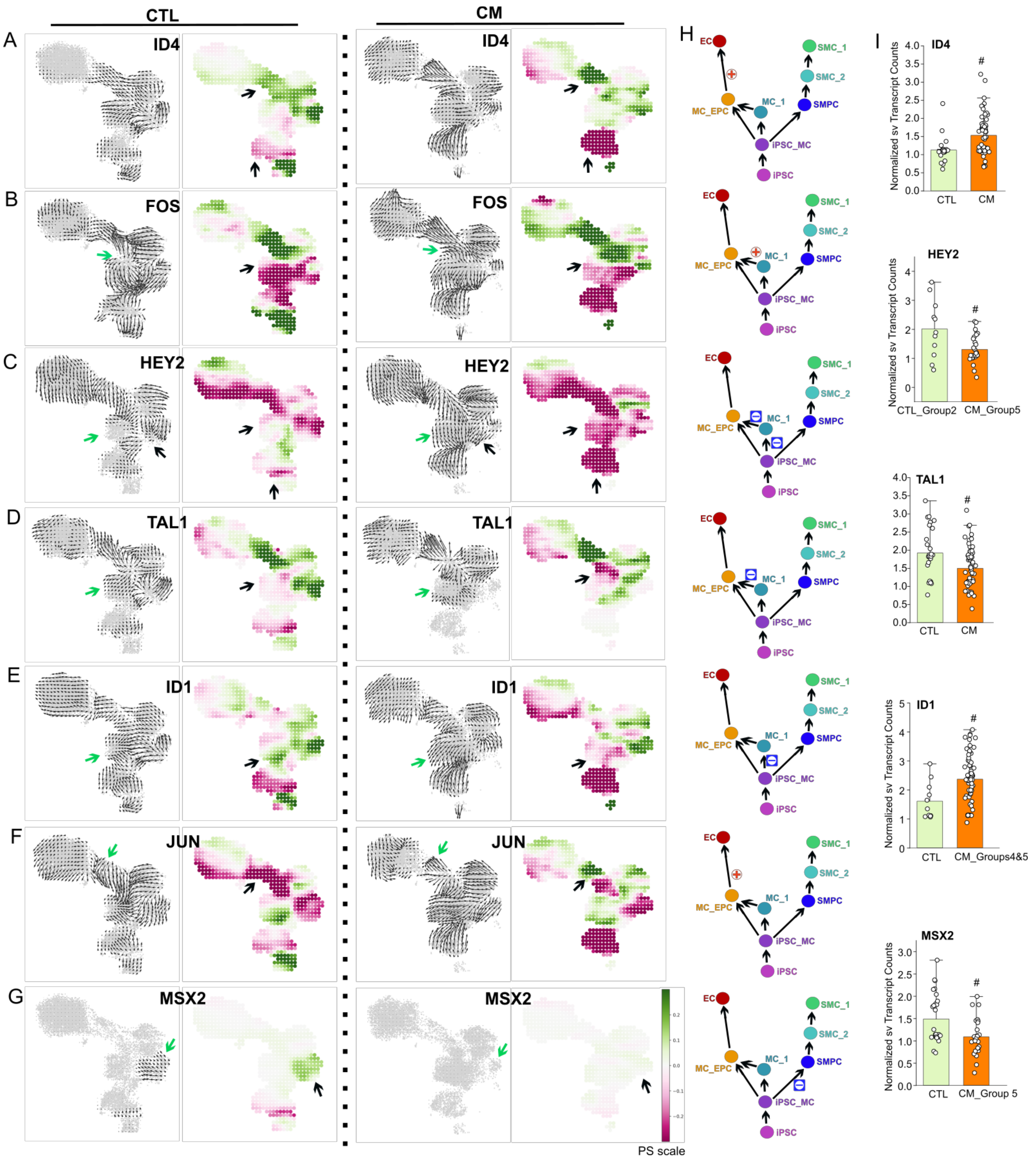


**Suppl figure 25** CellOracle simulation of cell-state transition in *Id4* (A), *Fos* (B), *Hey2* (C), *Tal1* (D), *Id1* (E), *Jun* (F), and *Msx2* (G) KO simulation in CTL and CM. Summarized simulation vector field and the perturbation scores were shown, respectively. PS scale, green color indicates differentiation promoted; red color, differentiation suppressed with specific TF KO simulation. H, Schematic of impact on perturbative shifts for iEC or iSMC lineages with each specific TF KO simulation in CM as compared to CTL. I, Single vessel GeoMx profiles of specific TFs in CM as compared to CTL. #, FDR < 0.05.

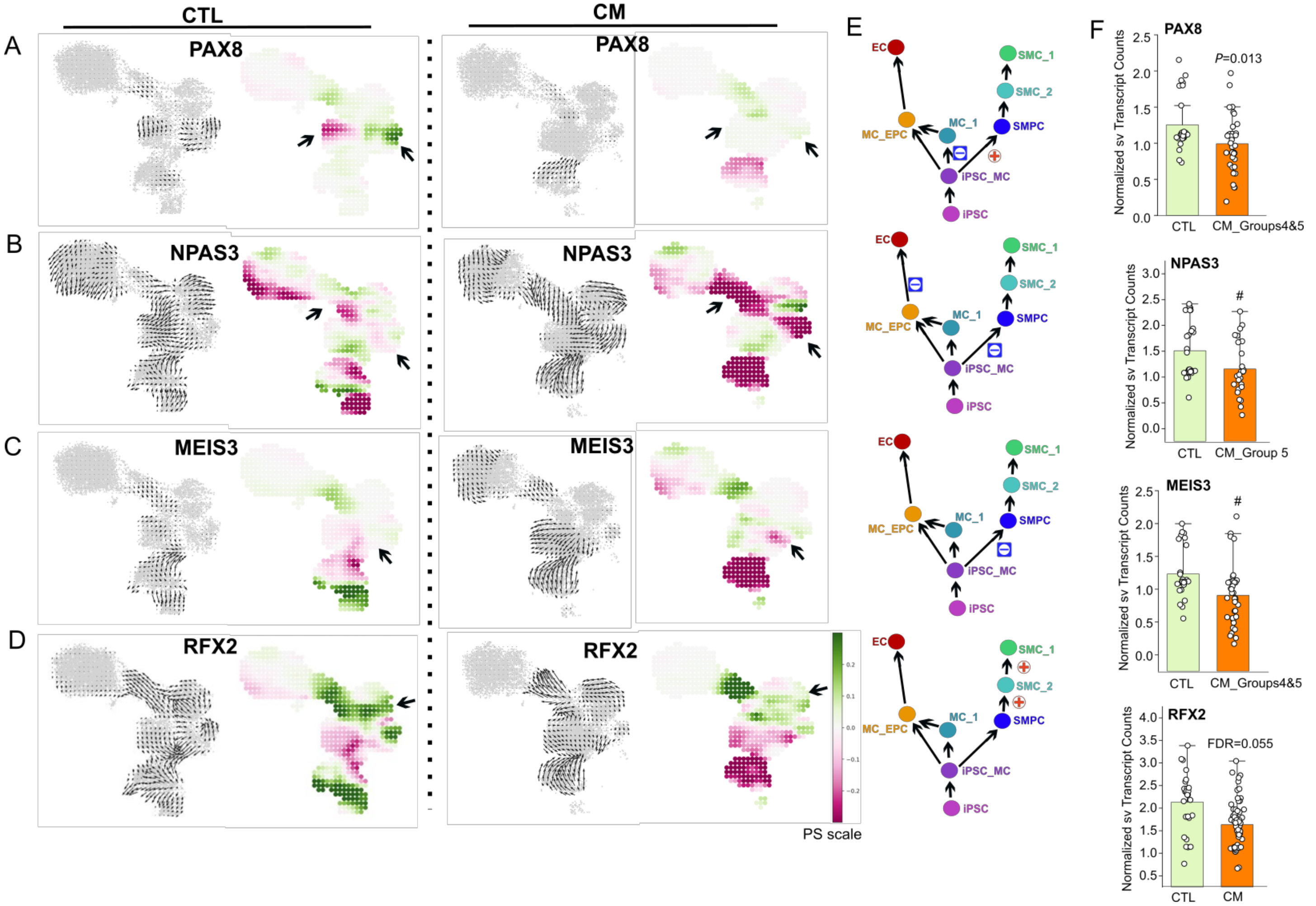


**Suppl figure 26** CellOracle simulation of cell-state transition in *Pax8* (A), *Npas3* (B), *Meis3* (C), and *Rfx2* (D) KO simulation in CTL and CM. Summarized simulation vector field and the perturbation scores were shown, respectively. PS scale, green color indicates differentiation promoted; red color, differentiation suppressed with specific TF KO simulation. E, Schematic of impact on perturbative shifts for iEC or iSMC lineages with each specific TF KO simulation in CM as compared to CTL. F, Single vessel GeoMx profiles of specific TFs in CM as compared to CTL. #, FDR < 0.05.
